# Spatiotemporal variation in size-dependent growth rates in small isolated populations of Arctic charr (*Salvelinus alpinus*)

**DOI:** 10.1101/2024.09.26.615117

**Authors:** Elizabeth A. Mittell, Camille A. Leblanc, Bjarni K. Kristjánsson, Moira M. Ferguson, Katja Räsänen, Michael B. Morrissey

## Abstract

As a key life-history trait, growth rates are often used to measure individual performance and to inform parameters in demographic models. Furthermore, intra-specific trait variation generates diversity in nature. Therefore, partitioning out and understanding drivers of spatiotemporal variation in growth rate is of fundamental interest in ecology and evolution. However, this has rarely been attempted due to the amount of individual-level data required through both time and space, and issues with missing data in important covariates. Here we implemented a Bayesian state-space model using individual-level data from 20 populations of Arctic charr (*Salvelinus alpinus*) across 15 capture occasions, which allowed us to: (1) integrate over the uncertainty of missing recapture records; (2) robustly estimate size-dependence; and (3) include a covariate (water temperature) that contained missing data. Interestingly, although there was substantial spatial, temporal, and spatiotemporal variation in growth rate, this was only weakly associated with variation in water temperature and almost entirely independent of size, suggesting that spatiotemporal variation in other environmental conditions affected individuals across sizes similarly. This fine-scale spatiotemporal variation emphasises the importance of local conditions and highlights the potential for spatiotemporal variation in a size-dependent life-history trait, even when environmental conditions are apparently very similar.

## Introduction

The size-dependent nature of many life-history traits, such as growth rates, and intra-specific variation within them, can impact diversity within populations, species, and ecological communities (1,2). Body size is interlinked with growth rate via various mechanisms, such as size-dependent feeding behaviour and metabolic rates (3). In fact, as body size is one of the most fundamental traits for organisms (4,5), influencing resource use (e.g., food choice; 6,7), competitive ability and reproductive success (e.g., larger individuals tend to obtain more mates and be more fecund; 8,9), and survival (10), it often covaries with individual fitness and life-history traits within populations, among populations and across taxa (11). As a life-history trait, growth rate is often used to assess individual performance (12) and variation in performance traits between individuals within and among populations influence population-level processes (13–15). In species with indeterminate growth, body size and growth rate can vary between individuals after maturation and can therefore continue to covary with fitness (14). Therefore, not accounting for individual variation explicitly in species with indeterminate growth, such as fishes, could be particularly problematic because unseen bias in predictions made from models using higher-levels of organisation can be increased (e.g., predictions of population dynamics based on growth rates; 16,17,18). Consequently, to robustly quantify variation in life-history traits and further our understanding of diversity in nature, we would benefit from accounting for size-dependence fully utilising individual-level data.

Despite the importance of understanding heterogeneity in organismal growth rates and the fact that the ecological and evolutionary processes that generate diversity vary in space, time and space-time (19–21), we have little understanding of spatiotemporal variation in size-dependent growth rates. Size-structured growth models that estimate the probability of transitioning between size classes have been around for a while (e.g., 22), but these models do not use individual-based data. Now that we know these models can lead to biased parameter estimates if there is heterogeneity among individuals in growth (17,which can be propagated into population models and erroneous conclusions drawn; 23,24), their use should be avoided when quantifying variation. The popular von Bertalanffy growth function (25) uses reasonable assumptions to relate growth rate to body size, and although it is hard to incorporate time-varying covariates into these models (e.g., those that change on time scales shorter than an individual’s life), it has recently been applied more widely to individual-level data (e.g., 16), including studies that investigate spatial variation in growth rates (26–28). However, even across taxa and traits, there are few datasets that are suitable for investigating spatiotemporal variation in growth rates due to the limited number of disconnected populations within a species that have been simultaneously sampled using mark-recapture through time (e.g., a few sites sampled within potentially connected populations or multiple distinct populations sampled for a few years, 26,27–29). Considering the lack of available data, it is unsurprising that we don’t understand whether temporal patterns of variation in traits such growth rates are spatially consistent or asynchronous. Therefore, an exploration of fine-scale spatiotemporal variation in growth rates using individual-level information could reveal interesting patterns.

Understanding heterogeneity in traits, such as growth, that may vary spatially (e.g., across populations) or temporally within generations, requires knowledge of environmental variation and repeated individual-level observations across time in multiple populations. In many species of ectotherms where environmental data is collected alongside organismal data, temperature covaries with growth rates (30,31). However, the direction and strength of covariation is not consistent across studies (32), indicating a context-dependent relationship between optimal temperature and growth (potentially due to an interaction between temperature and food availability; 33) that is not captured in current sampling schemes. The ideal of individual-based data are logistically demanding to collect in the wild (34). Therefore, most studies of growth rates and environmental covariates in wild populations have been conducted in single populations through time (longitudinal; 35), or in multiple populations at a single timepoint (27,cross-sectional; although see cases with a few populations; 36). Where individual-based data are available in multiple populations through time, the sampling sites may be connected within freshwater networks (i.e. not independent populations) and found across spatial scales where climatic conditions could also play a role (26,28). It is therefore not known to what extent increasing fine-scale replication in space and time will improve our knowledge about spatiotemporal variation. We can conceptualise how conducting either longitudinal *or* cross-sectional sampling in a natural system harbouring spatial and/or temporal variation in a focal trait (e.g., body size) could lead to an incomplete picture (Fig. 1) and result in inconsistent conclusions. For example, if there is spatiotemporal variation in a trait and a cross-sectional study is carried out (Fig. 1d), we may conclude that there is little trait variation within the system. If we imagine that quantifying variation in environmental covariates is affected in a similar manner, relationships between a focal trait and covariates may appear unimportant or not be considered at all. By collecting individual-level data and relevant environmental covariates, such as water temperature for ectotherms, across multiple populations within a limited area over time, questions about spatial and temporal variation in growth rates can be answered.

**Figure 1.**
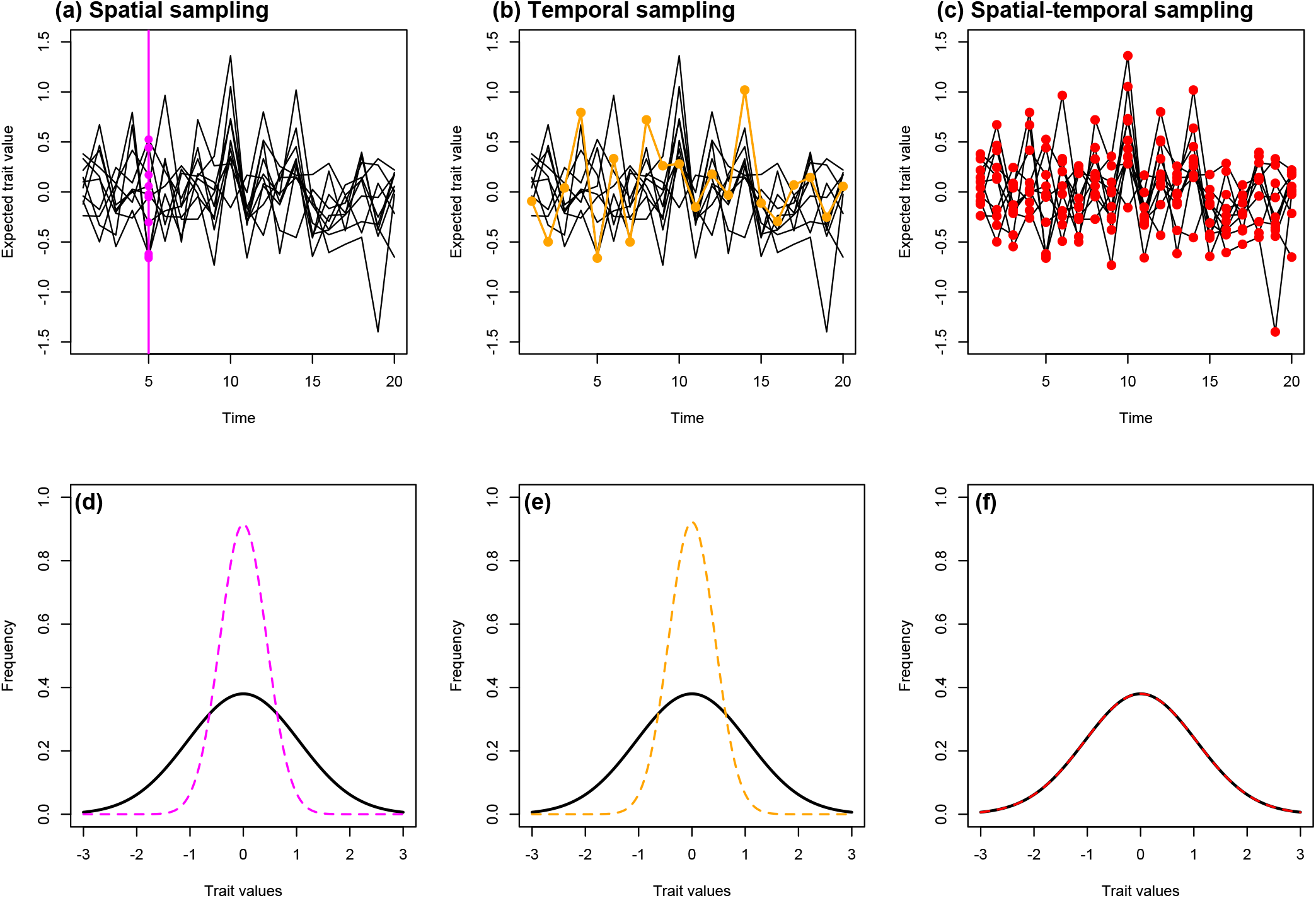
An example of variation in expected trait values at multiple locations (e.g., populations) of a meta-group (e.g., species) through time captured by different sampling schemes. The upper row shows the trajectories (lines) of the expected trait values that contain spatial-temporal variation. The vertical magenta line and dots represent a cross-sectional sampling scheme of all locations at a single point in time (a), which would obtain the spatial variation in trait values at that time point (d; magenta dashed line). The orange line and dots represent a longitudinal sampling scheme that follows a single location through time (b), which would obtain the temporal variation in trait values for that population (e; orange dashed line). The total variation of trait values across the meta-group are shown in black (d, e and f) and would be obtained in a sampling scheme that combined both cross-sectional and longitudinal sampling (red dots and dashed line in c and f).

Here we study growth rates of Arctic charr (*Salvelinus alpinus*) using highly spatially and temporally repeated sampling of individuals in a sub-Arctic system (37) where both size-dependence and temperature could be expected to influence spatiotemporal variation in intra- and inter-individual growth rates. In strongly seasonal environments, such as those found in sub-Arctic regions, long periods of resource limitation during winter months and temperature dependent processes (e.g., metabolic demands) would be predicted to differentially affect the growth rate of different sized individuals (3). This is because larger fish tend to have higher metabolic demands than smaller fish and so may be more resource limited during the winter period, but they can also survive starvation longer because the ratio of energy reserves relative to metabolic rate increases with size (3). Missing data is common in ecological studies and we explicitly demonstrate how to include and use information on an environmental covariate with missing data (in this case temperature), without removing or explicitly imputing data. We note that these complex size-dependent relationships between environmental variables and individual growth rate are not limited to fishes (examples range from phytoplankton to trees; 38,39), and so our methods have broad relevance. Overall, we present methods that are transferable across systems to investigate spatiotemporal variation in growth rates in populations where complex size-dependent processes may be occurring, and therefore, interesting results found.

In the current study, we used a unique dataset that contained 9247 body size observations of 3804 known individuals across 15 capture occasions from 20 small isolated populations of Arctic charr living in lava caves that were individually tagged as part of a long-term capture-mark-recapture (CMR) project. These populations are found within a small area (∼ 4 km^2^) in isolated pools within caves, fed by groundwater. Based on mark-recapture and population genetic analyses, these caves have limited connections (40), and therefore provide independent spatial repeats (as close to replication as can be achieved without manipulations in the wild). We had the following objectives. First we use individual-level body size data to ask how much variation is there in size-independent and size-dependent growth in terms of space (population), time (season and years) and space-time. Secondly, we ask how much more information do we obtain about the variation in growth rates by utilising these spatially and temporally replicated data compared to scenarios with fewer populations or shorter time-series; e.g., how big a component of the total variation in growth is temporal and could it be captured in a single population alone? Finally, we ask how much of the spatiotemporal variation in growth rate is associated with spatiotemporal variation in water temperature.

## Materials and methods

### Study system

Lake Mývatn in north-east Iceland (65°36’ N, 17°00 W) was formed after a major volcanic eruption around 2300 years ago (41,42). In the lava fields surrounding the lake there are numerous cave like formations that contain ponds fed by groundwater; these are particularly numerous in the Haganes and Vindbelgur areas (Fig. 2). Mývatn means “midge lake”, which reflects the large volume of midges emerging from the lake twice a year (42,43). These midges provide an important source of food for ecosystems surrounding the lake by falling onto the nearby land and into the caves (42,44).

**Figure 2.**
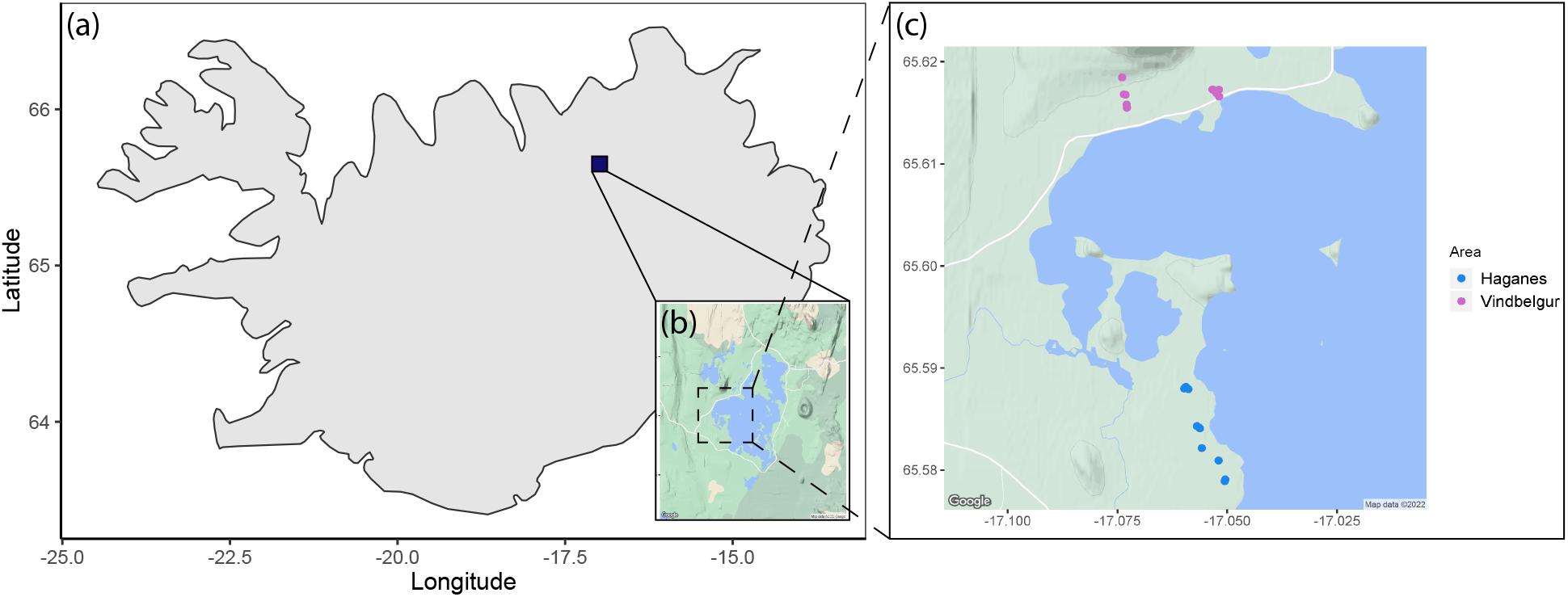
Map of the caves around Lake Mývatn in north-east Iceland. The maps show: (a) the location of the lake; (b) the general area of the caves; and (c) the finer-scale cave areas in Haganes (11 caves; blue) and Vindbelgur (9 caves; purple) within an ∼4 km^2^ area. Maps (b) and (c) are taken from Google Maps: accessed 29th July 2022.

Arctic charr are the only fish species present in most of these caves (Leblanc personal obs.) and are morphologically similar to the small benthic phenotype of charr (described in 45). The caves vary in the number and size of openings, as well as in their distance to the lake, and therefore in the potential amount of external food input (Leblanc *et al*., unpublished data). Individually tagged Arctic charr from 20 caves have been monitored since August 2012 (37). These study caves range in size from very small, where only a few fish are caught at each capture occasion, to relatively large, where over 100 fish can be caught at a time. The full extent of underground connections among caves is unknown. However, analyses of the population genetic structure and inter-cave recaptures (or lack thereof) suggest that these 20 study caves consist of 16 unique populations (40), indicating that only a limited number of caves are connected.

### Data collection

To obtain data on Arctic charr growth rates and fine-scale environmental variation, the 20 caves were visited in June and August over eight years (2012 to 2019). At each visit fish were trapped, individually marked and their body size measured. Each month/year combination is referred to as a capture occasion. During each capture occasion, two visits were made to each cave one to two weeks apart to increase the capture probability of fish. These sampling months approximately coincide with the growing season of Arctic charr, from the beginning of the Icelandic summer (June) to the near the end (August). Although it is likely that the growing season starts in May, continues some weeks after August and varies year to year, this sampling period should capture the bulk of summer growth across years (i.e. coinciding with food peaks; 42,43). Therefore, we refer to the period between June – August as “summer” (∼ two months) and the period between August – June as “winter” (∼ 10 months).

Fish were trapped using minnow traps and electrofishing. Upon capture, body size was measured (fork length, nearest mm) and a photograph was taken of their left flank (37). In June 2019 repeat measures of fork length were made on a random subset of 76 individuals in the field to estimate the measurement error variance, which should capture the majority of the technical variance in measurement error. Upon first capture, fish were tagged and the upper lobe of the caudal fin clipped to obtain a tissue sample for later genetic analyses. Fish with fork lengths ≥ 65 mm were tagged with passive integrated transponder (PIT) tags (12 mm HDX; Oregon RFID), whereas fish with fork lengths between 45 – 64 mm were tagged using either Visible Implant Elastomer tags (prior to June 2015) or PIT tags (from June 2015 onwards; 8 mm HDX; Oregon RFID).

We observed some instances of lost PIT tags, as evident from individuals with no PIT tag but with visible scars or fin clips from sampling for genetic analysis (Leblanc personal obs.). Although the incidence of tag loss is low, the frequency is unknown. Efforts to re-identify fish using photographs, either manually or using computer deep-learning methods, suggest tag loss rates of approximately 5–10% (e.g., 6.5% and 7.5% in two caves over the entire study; 37). For the purposes of the analyses of growth in this study, we expect tag loss to result in a reduction in the completeness of individual recapture records, but we do not expect it to cause systematic errors in the partitioning of variation in size-dependent growth rates.

Given that Arctic charr is an ectotherm, we were particularly interested in the association between individual growth rates and water temperature. A temperature logger (UA-001-64 Pendant temperature HOBO; Onset Corporation) was used to measure water temperature in each of the 20 caves four times daily throughout the year from 2013 onwards. The average water temperature in each cave was estimated for each *k* using January temperatures for the “winter” period and July temperatures for the “summer” period. A small amount of missing data exists in this dataset as not all *cave* × *k* combinations had temperatures recorded during these months. Out of the 300 *cave × k* combinations, there are 234 with an estimate of water temperature. Overall, there are 9247 fork length observations from 3804 individual fish; of these, 7046 are associated with an observed water temperature measurement and 2201 are not.

### Analyses

#### General overview of the approach

Not every individual was measured at every capture occasion, which could be due to either death or a failure to capture. Therefore we used an integrated state-space model containing linked process models for growth and time-/space-dependent temperature, and, likelihoods for time series data on individual size and temperature. This model is set-up in a Bayesian framework to allow for missing information in individual growth trajectories without the need for either imputation or subsetting to a complete dataset (46). In this method, observed data are utilised during parameter estimation of data-poor individuals by ‘borrowing strength’ from observations of similar individuals (47). This allowed us to use fork length measurements from full capture histories by including predicted fork lengths (with full integration over uncertainty) when individuals were not captured; there are other methods of prediction (e.g., empirical Bayes and maximum likelihood; 27,28), but these do not fully integrate over the uncertainty in body size. To estimate growth rates based on fork lengths of Arctic charr within the lava caves, a Bayesian state-space model was implemented in R version 3.5.1 (48) using the ‘rjags’ package (49). For both growth and water temperature, observation models informed process models. The variation in the parameters of the process models provided size-independent and size-dependent estimates of spatial, temporal and spatial-temporal variation in growth rates, spatiotemporal variation in water temperature and spatiotemporal variation in water temperature that is associated with spatiotemporal variation in growth rate. In our study spatial variation refers to the variation among caves averaged across years, temporal variation is the variation among years averaged across caves, and spatiotemporal variation is the additional variation which is not accounted for by cave or year. All estimates are reported as posterior means with 95% credible intervals [CI] in brackets.

The model used here is a form of linear mixed-effects model. It has been shown that the variance estimates from these types of models are robust to violations of distributional assumptions (50), and avoid problems arising from pseudo-replication (e.g., fish live in the same environment between capture events; 51). Importantly, our approach differs from other models in the implementation of a data augmentation scheme for both modelling body size on occasions when an individual was not captured *and* a covariate with missing data, which allows us to maintain our sample size whilst avoiding data imputation. Although others have used latent state models for body size in fish (e.g., 52), they do not consider using the same approach for covariates. We show that this is possible and outline the limits of the method, which should inform future work on how to obtain the most information possible from datasets. As part of this, we attempted to include variation in the input of terrestrial resources into the caves as a covariate in our growth model. However, we found that the lack of spatial and temporal replication of these data caused numerical issues that meant the model could not converge. We therefore only focused on water temperature as a key environmental proxy for ectotherms.

We carried out model validation and used posterior predictive checks to see how well the model was able to predict fish size when it had not been measured. To generate the data for this we randomly removed a single observation from ten randomly chosen individuals and re-ran the model; a process that we carried out 99 times to obtain 990 predicted values for comparison with the observed data. All models converged well and passed these checks, with: ∼ 93% of the observed data falling within the 95% CI of predicted values, a Pearson’s correlation coefficient between the predicted and observed values of *r* = 0.982 and a mean absolute residual (observed - predicted) of 2.85 mm (see Supplementary Information for full methods and results).

#### Temperature model

The temperature data contained missing values with 22% of *cave* x *capture* combinations missing observed data. Covariates cannot contain missing data. Therefore, in order to incorporate the most plausible information for this covariate into the growth model, we used data augmentation (for detailed information on data augmentation see 46). The temperature process model explicitly modelled the two seasons (*a*_*h*_) and allowed varying intercepts across caves (*a*_*hj*_) and years (*a*_*ht*_)

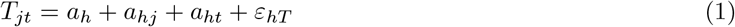

where *T* refers to temperature and *ε*_*hT*_ is the residual variation. This is the additional variation not accounted for by the effects of season, season-cave, or season-year. A temperature observation model informed the temperature process model

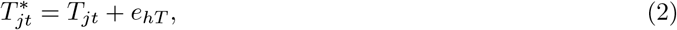

with 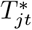 referring to the observed temperature for each *cave × capture* combination. The error variance for each season is essentially zero but was included to enable the data augmentation.

#### Growth process model

In the growth model, the size (*z*) of an individual *i*, in cave *j* at time *t* + 1, was dependent on the size of individual *i* at *t*

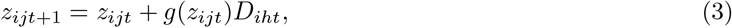

where *g*(*z*_*ijt*_) is the growth function and *D*_*iht*_ is the number of days since an individual was last captured (scaled by season). For fish that were not caught on a consecutive capture occasions the average mid-date of the two visits across years was used to calculate *D*_*iht*_ (16th June and 22nd August). This accounted for any differences in the time of capture on estimates of growth rate. Growth was modelled

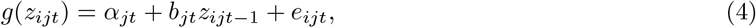

where *e*_*ijt*_ are residuals. Although, size-at-age functions are nonlinear, their derivatives (e.g., the von Bertalanffy growth function; 25) are linear, and therefore it is typical to fit a linear model to growth (e.g., 16). Varying slopes (i.e. size-dependent growth; *b*_*jt*_) and intercepts (i.e. size-independent growth; *α*_*jt*_) were allowed for the spatial (*j*), temporal (*t*) and spatial-temporal (*jt*) components

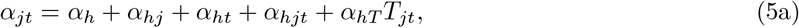

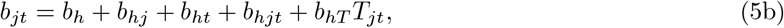

where *h* indicates the season (summer or winter). Explicit modelling of the two seasons allowed us to partition spatial and temporal variation into what are expected to be a rapid growth season (summer) and a slow growth/maintenance season (winter) for these populations. The spatial, temporal and spatio-temporal components for both intercepts and slopes were modelled as random effects, drawn from a multivariate-normal distribution, with estimated variance and covariance. It is from here that the desired variances are obtained, for example, *α*_*j*_ ∼ *N* (0, *V*_*αj*_), where *V*_*αj*_ is the spatial variation in size-independent growth. A normal distribution was assumed for the size of individuals at first capture, 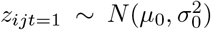 and uniformative priors were used (see code for the full models and priors). Water temperature is explicitly included as an effect in the parameterisation of size-dependent growth (Equation 5b; indicated by *T*)3. Finally, all models used mean-standardised data (i.e. centring using mean size and water temperature).

#### Growth observation model

The growth process model was informed by an observation model using the observed fork lengths

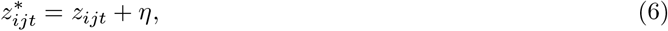

Where 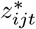 is the observed size of an individual *i*, in cave *j* at time *t*. The measurement error variance, *η*, was estimated using the repeat measures made in June 2019 in an intercept model with individual fish identity as a random effect 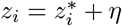, where 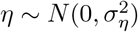. In addition to the main model outlined here, models were run without temperature and season effects independently to test whether season absorbed all the variance in growth rates driven by water temperature. This was not the case (see Supplementary).

#### Variance component derivation and interpretation

Examples of the notation used for variance components here are: 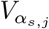 which represents spatial variance (*j*) in the size-independent (intercept; *α*) growth rate in summer (*h* = *s*); and 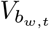 which represents temporal variance (*t*) in the size-dependent (slope; *b*) growth rate in winter (*h* = *w*).

The variances in growth rate related to variation in water temperature was estimated using the mathematical rule *V* (*a* + *bx*) = *b*^2^*V* (*x*) (as in 53). For example, temporal variation in size-independent growth related to variation in water temperature in summer 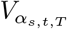 was estimated,

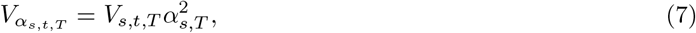

where *V*_*s,t,T*_ is the temporal variation in water temperatures (i.e. variance of the random effect of year on water temperature) in summer and 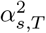 is the effect of temperature on size-independent growth taken from Equation 5a.

The variance in size-dependent growth rates (slopes; 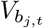) is not easily comparable to the variance in size-independent growth rates (intercepts; 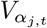). For example, small values of 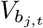 may not represent small values of 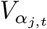 when they are on the same scale. Therefore, we calculated the amount of variability in 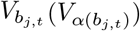 that equated to variation in the intercept from the product of the variance component of interest and the variation in the corresponding data. For example, the variability in spatial variation in size-dependent growth rates in summer 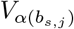 is calculated as

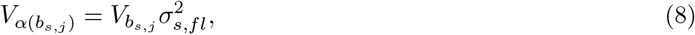

where 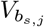 is the spatial variation in size-dependent growth rate in summer and 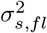 is the variation in the fork lengths of fish after a period of summer growth. As the growth model does not explicitly obtain the variation in the fork lengths 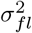, these were taken from mixed-effects models run in MCMCglmm (54) for summer and winter separately (see Supplementary).

To assess how representative the size-dependent and size-independent variation in growth is over space, time and space-time, the proportion of the spatiotemporal variation in growth (*R*) that each component made up was calculated. For example, the amount of total variation in growth that the spatial variation in size-independent growth 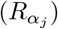 represents was estimated as

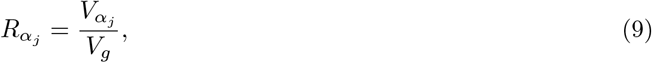

where 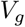 is

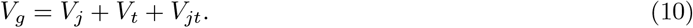

Each variance component here, for example *V*_*j*_, contains the size-independent 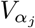, the size-dependent 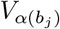 and the temperature associated 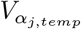 growth variances in the calculation of *V*_*g*_. Both Equation 9 and 10 were calculated separately for summer and winter.

Finally, to visualise spatiotemporal variation in growth rates in terms of the relationship between size and growth (i.e. size-dependent growth), growth functions of fish in the core size range for these fish (70-120 mm) were simulated using the parameter estimates from the model for summer and winter separately.

## Results

### Water temperature

The average water temperature in the caves was 6.33 °C [5.84; 6.79] in summer and 4.32 °C [3.81; 4.73] in winter (Table 2). There was more variation in water temperature in winter than in summer, mainly due to a larger source of unknown variation (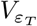; Table 2). In both seasons, variation in water temperature among caves was greater than variation among years. The between year variation in water temperature was relatively low in winter compared to summer.

**Table 1:** Spatial and temporal variation in cave water temperature estimated from the model for summer and winter separately. The posterior mean is the overall mean across all caves and years for each season. The 95% credible intervals are shown in brackets.

|  | *Summer* | *Winter* |
| --- | --- | --- |
| Posterior mean ($\bar{T}$) | 6.33 [5.84; 6.79] | 4.32 [3.81; 4.73] |
| Variance among caves ($V_{a_j}$) | 0.22 [0.02; 0.82] | 0.21 [1.31e-03; 1.10] |
| Variance among years ($V_{a_t}$) | 0.15 [0.02; 0.51] | 0.03 [6.64e-04; 0.19] |
| Residual variance ($V_{\varepsilon_T}$) | 0.35 [0.26; 0.46] | 1.03 [0.77; 1.36] |
| Total variance ($V_T$) | 0.72 [0.30; 1.79] | 1.27 [0.77; 2.00] |

**Table 2:** Variance components associated with growth through time, space and space-time shown for summer and winter separately, including size-independent, size-dependent and temperature-associated growth. The variability in growth associated with the size-dependent variance components is shown. *R* is the proportion of the variation in growth rates that spatial and temporal variation account for. Posterior means are reported and 95% credible intervals are shown in brackets.

|  |  | *Summer* | *Winter* |
| --- | --- | --- | --- |
| Size-independent variance components | Space $V_{\alpha_j}$ | 1.986 [0.809; 4.490] | 5.032 [2.025; 11.521] |
| | Time $V_{\alpha_t}$ | 4.721 [1.130; 21.369] | 16.593 [4.759; 63.877] |
| | Space-time $V_{\alpha_{j,t}}$ | 1.233 [0.637; 2.098] | 5.901 [3.953; 8.596] |
| Size-dependent variance components | Space $V_{b_j}$ | 0.001 [2.81e-04; 0.002] | 0.001 [3.40e-04; 0.003] |
| | Time $V_{b_t}$ | 0.001 [2.07e-04; 0.003] | 0.001 [3.30e-04; 0.005] |
| | Space-time $V_{b_{j,t}}$ | 0.001 [2.92e-04; 0.001] | 0.002 [0.001; 0.003] |
| Variability in growth associated with size-dependent variance components | Space $V_{\alpha(b_j)}$ | 0.025 [0.006; 0.069] | 0.045 [0.010; 0.133] |
| | Time $V_{\alpha(b_t)}$ | 0.026 [0.004; 0.107] | 0.101 [0.014; 0.412] |
| | Space-time $V_{\alpha(b_{j,t})}$ | 0.437 [0.168; 0.854] | 0.962 [0.381; 1.785] |
| *Temperature-associated* variance components | Space $V_{j,T}$ | 0.114 [0.004; 0.715] | 0.000 [5.45e-06; 0.287] |
| | Time $V_{t,T}$ | 0.080 [0.003; 0.462] | 1.97e-06 [1.16e-06; 0.046] |
| | Space-time $V_{j,t,T}$ | 0.224 [0.015; 0.688] | 0.000 [1.70e-04; 0.913] |
| Size-independent variance components repeatability | Space $R_{\alpha_j}$ | 0.240 [0.060; 0.483] | 0.190 [0.049; 0.396] |
| | Time $R_{\alpha_t}$ | 0.489 [0.192; 0.843] | 0.539 [0.264; 0.845] |
| | Space-time $R_{\alpha_{j,t}}$ | 0.151 [0.042; 0.300] | 0.222 [0.073; 0.396] |
| Size-dependent variance components repeatability | Space $R_{b_j}$ | 1.02e-04 [2.12e-05; 2.73e-04] | 4.28e-05 [8.39e-06; 1.19e-04] |
| | Time $R_{b_t}$ | 9.94e-05 [1.77e-05; 3.52e-04] | 4.81e-05 [1.10e-05; 1.46e-04] |
| | Space-time $R_{b_{j,t}}$ | 9.06e-05 [2.15e-05; 2.07e-04] | 5.87e-05 [1.49e-05; 1.29e-04] |
| Residual variation | $V_e$ | 9.403 [5.701; 16.612] | 17.539 [8.249; 43.244] |
| Total variation | $V_g$ | 19.581 [12.018; 36.166] | 50.169 [27.618; 98.777] |

### Charr size and growth

The overall mean fork length was 92 mm [95% range 52-134 mm], and the average increase in fork length of a 92 mm fish was 3.4 mm/month [2.41; 4.46] in summer (total 6.79 mm [4.82; 8.92] over the two summer months), and 0.82 mm/month [0.42; 1.21] in winter (total 8.18 mm [4.25; 12.07] over the 10 winter months). Water temperature had a small effect on this average growth in the summer (0.40 mm/month [0.10; 0.69]), but no detectable effect in the winter (−9.85e-04 mm/month [-0.08; 0.08]). Overall, larger fish grew less than smaller fish (annual size-dependent slope estimate ≈ -0.12 [-0.19; -0.07]). This relationship between individual size and growth was greater in the summer (summer size-dependent slope estimate ≈ -0.05 [-0.06; -0.03]) compared to the winter (winter size-dependent slope estimate ≈ -0.02 [-0.02; -0.01]) on a per month basis. There was no detectable effect of water temperature in the two seasons with the estimated effects on the size-dependent slopes in summer: 4.99e-03 [-3.76e-03; 0.01]; and winter: 7.81e-04 [-9.21e-04; 2.48e-03].

Fish caught after the summer were larger on average (94.88 mm [90.07; 99.81]) than fish caught after the winter (90.28 mm [83.55; 97.51]). The measurement error variance of fork length was ∼ 0.6 mm^2^.

### Seasonal variance components

Total variance of growth rate in winter was double the variance in summer (*V*_*g*_; Table 3). Of this variance, a considerable proportion was partitioned as size-independent components of growth (intercepts), particularly in time and in the winter. In contrast, relatively less variation was partitioned as size-dependent estimates (slopes) in space, in time, and in the interaction of space and time, in both seasons. Note, however, that because the variance in slopes and variance in intercepts are in different units, these variances cannot be directly compared. Our calculation of the variability in growth rates arising from heterogeneity in the size-dependent growth parameter (see eq. 8), converts the variance into the associated variance in growth. The size-dependent values increased when the variances in slopes were expressed in terms of the associated variability in growth rates (i.e. in the same units as variance in intercepts; Table 3; 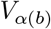. However, they still remained relatively small compared to the size-independent variance components (e.g., 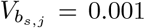 *mm* of growth per *mm*^2^ increase in fish size, equates to variability in growth associated with 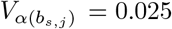 *mm*^2^, compared to 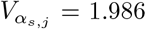 *mm*^2^). Additionally, we constructed a visualisation of variability in size-dependent growth functions, specifically to illustrate the variability arising from intercepts and slopes, by simulating random draws from the distributions of intercepts and slopes, and plotting them (Fig. 3). This allowed us to assess the consistence (or otherwise) of slopes, in relation to the degree of variability of intercepts (elevations of the functions) graphically. These simulations showed that individual growth across space, time and space-time was lower, less dependent on individual size and less variable during winter (Fig. 3d-f) compared to the summer (Fig. 3a-c).

**Figure 3.**
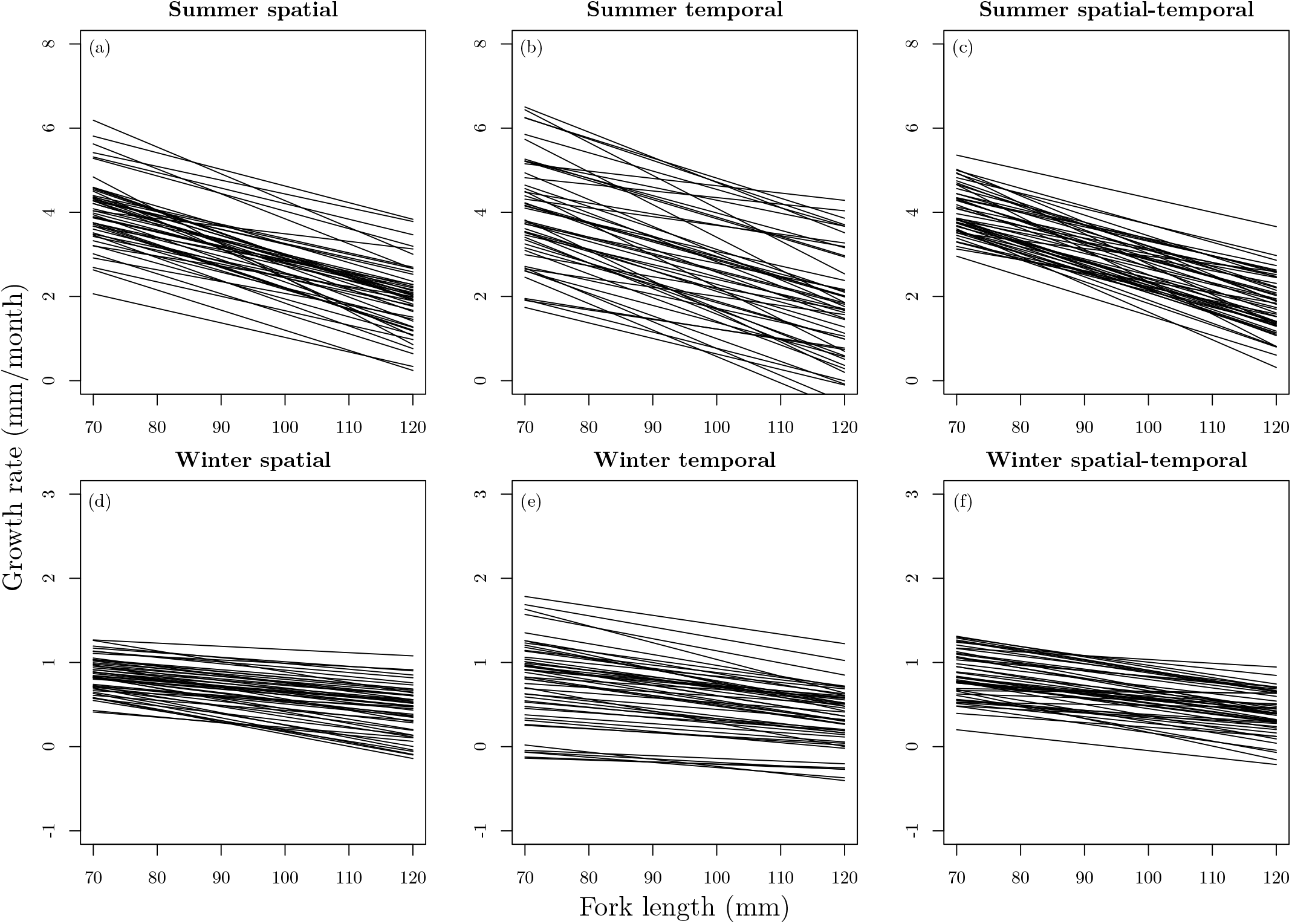
Simulated growth functions using fish sized 70 – 120 mm in summer (top row; a-c) and winter (bottom row; d-f) including spatial, temporal and spatial-temporal variation (left, middle and right plots, respectively). The x-axes show the size of the fish in mm and the y-axes show the growth rate in mm/month. Note that the scale of the growth rate in summer and winter differ.

Size-independent spatial, temporal and spatial-temporal variation were all substantial proportions of total variation in growth (*R*_*α*_; Table 3). Notably, temporal variation in size-independent growth accounted for approximately half of the total variation in growth, in both summer and winter. Calculating the amount size-dependent variance components represented in total variation in growth resulted in very small values because the variation in size-dependent growth was so small.

The trend in the temperature-associated variance components differed between the two seasons, with very little variation in the winter and more space-time than space or time variation in the summer (*V*_*T*_ ; Table 3). However, overall variation in water temperatures and variation in growth rates were only weakly associated. This is visualised in the lack of trend between the spatial-temporal variation of size-independent growth, those with the largest variation, and the spatial-temporal variation in water temperatures in summer and winter (Fig. S5).

## Discussion

Using a unique dataset containing individual-level longitudinal data in multiple small isolated populations of Arctic charr that inhabit lava caves around Lake Mývatn, Iceland, we were able to partition spatial, temporal and spatial-temporal variation in growth rates. Much of the variation in growth rates was size-independent rather than size-dependent, suggesting that good or bad environmental conditions tended to affect fish of all sizes similarly. As with most capture-mark-recapture studies, missing data was present within our heterogeneous individual growth trajectories, which can be an issue using frequentist analyses (35). Here we implemented Bayesian state-space modelling which enabled us to investigate variation within and among caves over time using data containing missing values. Although this type of modelling has been successfully implemented in other studies (35), the degree of replication used here is high relative to other space and/or time studies in natural populations. Our partitioning of the spatial component alongside the temporal component of variation in life history traits emphasizes the importance of local conditions in these (mostly) isolated populations. This is of interest for both evolutionary ecology and conservation biology as, for example, this type of information is used to inform parameters in demographic models and/or management decisions (55,56), which is only possible with temporally consistent data across multiple populations.

This dataset also allowed us to demonstrate that a significant proportion of the variation in growth rate of Arctic charr in these caves would not have been captured with lower spatial and temporal replication (Fig. 1; *R*_*α*_; Table 3). For example, by having spatial repeats that allow explicit partitioning of spatiotemporal variation, we can be more confident that the temporal variation found is consistent across caves because any space-time specific variation is partitioned out separately. This emphasises the importance of simultaneous long-term individual-based data collection and ecological monitoring in multiple locations. Finally, we showed that it is possible to include an environmental covariate with missing data, in this case water temperature, in the models. Doing this we found that water temperature accounted for a relatively small amount of the variation in growth rates. Although this might be unexpected as temperature is thought to impact ectotherm growth ubiquitously (Angiletta 2009), it makes sense here because there is little variation in the water temperature itself (95% range: 2.73 – 7.38 °C). Overall, despite stability in an apparently important driver of variation, our work highlights the potential for spatiotemporal variation in a size-dependent life-history trait, and therefore, should not be overlooked.

### Spatiotemporal variation in growth rates

We found little spatiotemporal *variation* in size-dependent growth (*V*_*b*_; Table 3), even though individual growth was size-dependent (i.e. larger fish grew less than smaller fish; size-dependent slope estimate ≈ -0.12). Fish growth is often size-dependent (25) due to ontogeny (57) and size-structured competition (58). For example, younger and smaller fish typically grow faster than, and are prey for, older larger fish (the ontogenic component; 57). Likewise, dominant (probably larger) fish within cohorts often grow faster than subordinates as size-structured dominance hierarchies within populations create inequalities between individuals in feeding opportunities (58). However, the little *variation* in size-dependent growth in these Arctic charr suggests that variation in environmental conditions affected fish in the caves equally regardless of their size and that size-structured competition could be relatively stable in space and time.

The lack of variation in size-dependent growth in these Arctic charr could be due to equally successful opportunistic feeding across size classes in these thermally stable cave environments (i.e. little variation in water temperature; Table 2). These lava caves are part of a young ecosystem with relatively low productivity (42) and the fish inhabiting them could be limited across all sizes. That is, even in the case where larger individuals are out-competing smaller individuals in terms of resource use, individuals of all sizes may be merely meeting their metabolic demands rather than thriving during resource scarce periods (e.g., as found in other Arctic charr populations; 3). Although some prey items may be available in the benthic zone within the caves for most of the year, during the short-lived midge emergence peaks that occur in the Mývatn area (typically twice a year in spring/summer; 43) there is a large influx of terrestrial prey onto the surface water at the cave openings (Leblanc personal obs.). These midge and other invertebrates entering via cave openings present a different resource for the Arctic charr to feed upon. Individuals in other populations of small benthic Arctic charr across Iceland have been found to have a diverse diet, often including terrestrial prey (45), indicating generalist opportunistic feeding. Furthermore, the energy expenditure for these fish is likely to be low due to limited predation and water flow, which could be one reason that these fish are able to grow in these low water temperatures. Overall, the combination of opportunistic feeding, metabolic rates that vary with size and the constant low resource conditions in the cave environment could have led to consistent growth rates across sizes (and potentially adaptation; 59), seen as a lack of variation in size-dependent growth here.

The considerable temporal variation in size-independent growth observed in our study, particularly during winter, may be due to both variation in terrestrial food sources and the timing of our sampling. The abundance of epibenthic chironomids and Cladocera vary temporally within Lake Mývatn itself (42), with one chironomid species (*Tanytarsus gracilentus*) in particular showing dramatic temporal fluctuations in its population abundance (44). Although other salmonids are adapted to large and predictable, but short-lived, resource pulses via physiological changes to their digestive machinery (e.g., the digestive tract is allowed to atrophy during periods of resource scarcity; 60), among year temporal variation in resources of our Arctic charr system may be too unpredictable for similar adaptations to be present. Instead, the temporal variation of invertebrate abundance in Lake Mývatn among years might be mirrored in the caves with the terrestrial and aerial input of invertebrates (44) into the caves varying through time, which could explain some of the temporal variation in size-independent growth. In addition, some of the temporal variation associated with the winter months could be a result of variation in the timing of the winter period relative to its definition in our study: due to the differing lengths of the two seasons, winter growth rates were less variable per month than summer growth rates (Fig. 3). Specifically, although the seasonal transition in the Mývatn area is known to vary year-to-year (42), the two seasons used here were defined by our fixed sampling scheme (i.e. similar timing each June and each August). Therefore, annually varying lengths of the growing season (e.g., early snow melt, warm temperature years) are not captured by our sampling schedule and could be part of the temporal variation we found in size-independent growth. Overall, temporal variation in size-independent growth suggests that ecological conditions in the caves vary across years, with total variation being greater in winter months, but variation on a per month basis being greater in the summer.

The spatial variation seen in size-independent growth in this system (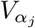; Table 3) could be the result of variation in the size and orientation of the caves, the size of their openings and their distance to Lake Mývatn. This physical variation among the caves is likely to contribute to variation in the opportunity for primary production, the zoobenthos within them and the amount of input from the surrounding terrestrial environment. For example, the midges that emerge from Lake Mývatn rapidly decline in abundance with distance from the lake, which occurs to a greater degree during the weeks of peak midge emergence (44). Therefore, the influence of the physical features of the cave are one possible source of spatial variation in growth rates due to potentially concurrent resource variation.

### Capturing spatiotemporal variation in growth rates

It is generally difficult to study spatial and temporal variation simultaneously, and include within-population variation, but we have shown here that such variation is more than a sum of its parts. Importantly, a significant proportion of the variation in growth rates of these charr would not have been captured without the spatial and temporal replication (Fig. 1; *R*_*α*_; Table 3). For example, the temporal variation would have been missed in a cross-sectional study, lowering the estimates of variation in growth rate. This can be crucial as the parameters obtained from these types of models can feed directly into population-level models such as IPMs and influence what covariates we might consider to be important for a focal trait. Furthermore, the considerable individual variation in the growth trajectories of fish in this study system (examples in Fig. S1; *V*_*e*_; Table 3), highlights the importance of including within-population variation in our models rather than using population trait means. This is something that is often disregarded in ecological models (reviewed in 61). If we had ignored this individual variation, our estimates of variation in growth rates would have likely been upwardly biased (17), leading to false conclusions. For example, larger between-individual variation within some caves than others could have inflated our estimates of spatial variation in growth, and we might have concluded that cave-effects were stronger than they are in this system. However, by explicitly including this within-population variation in our model, we avoided these issues and captured robust estimates of spatial and temporal variation in growth. Therefore, our results emphasise the importance of individual-based studies being carried out in time and space.

### Variation in growth rates associated with variation in water temperature

Although fish can be sensitive to temperature changes of just 0.5 °C (62), most of the variation in growth rate was not explained by water temperature in the caves in our study, which could partly be explained by the caves thermal stability. In general, the average water temperatures found in these caves were comparable to, but more stable than, many of the freshwater systems Arctic charr inhabit (3,Table 2; 63). In both seasons, the variation in water temperature among caves was greater than the variation among years, but there was relatively little variability overall. The stability of water temperatures within the caves through time and the consistent spatial variation seen among caves could be explained by the thermal stability that characterises cave ecosystems (64) and groundwater sources further stabilising water temperatures (42,65). The small amount of temperature-associated variation in growth rates (*V*_*T*_ ; Table 3) could be driven by temporal variation in summer water temperatures and caves found in consistently warmer areas. As ectotherms, the energy costs and gains of salmonids are affected by water temperature in a size-dependent manner, which explains why water temperature is often an important driver of variation in salmonid growth rates (e.g., 63,66), but this is not always the case (18,e.g., 67,68). Optimum growth rates of Arctic charr are thought to occur in water temperatures of 12-14 °C in the wild, in fish from wild populations reared in common garden set-ups, and in aquaculture strains (67,69). Interestingly, the temperatures found in our study caves were consistently between 4 to 7 °C, which would appear to be below their optima (although we note that there can be an interaction between temperature and food availability in determining the optimum; 33). Hence, it could be that these fish are never growing at their full potential [approximations of the asymptotic size of these charr ranges between ∼ 100 − 266 mm; Supplementary Table 4; in part explaining their small size relative to many freshwater charr which can reach 71 cm in length; (70)], and/or have adapted to a colder environment (45). Overall, water temperature explained little of the variation in growth rate, which probably reflects the stability of the water temperature in this distinct system.

### Wider applicability of the method

As individual-based CMR data is always likely to contain missing data in both individual information and covariates, one direct application of the method used in this study would be to improve the estimation of additional key life-history parameters, such as survival (see similar work in single populations for reproductive output; 71). The data augmentation scheme implemented here via Bayesian state-space modelling allowed us to integrate over the size each individual was when it was not seen and include an environmental covariate with missing data. Whilst we were able to account for missing data, we note that there is a limit to the amount of missing data that can be present in covariates. The 78% coverage across both seasons for water temperature was sufficient, whereas the 60% coverage within summer alone (30% across both seasons) for terrestrial input was not (models did not converge and so there are no results to present). This highlights that there is a limit to the missing data in the predictor variables that can be used within these models, which would need to be taken into account in the application of this framework to ensure that the number of spatial and temporal repeats were sufficient for partitioning the variation in the focal traits. Therefore, in order to carry-out such analyses a non-trivial dataset is required, but when data is available this is a powerful approach to use to maximise the information contained in such datasets. Furthermore, it is useful to be aware of what information might be lost under different study designs (Fig. 1), both prior to design and once the data has been collected. Overall, we have demonstrated that these models can integrate over the uncertainty of missing data within individual growth trajectories and a covariate to partition spatial and temporal variation in an important life-history trait, an approach that could be applied to other systems.

## Conclusion

Not only have we noted the limits of data required to include covariates with missing data, but we have also demonstrated how trait-dependence can be incorporated into individual-based models to partition out variation in time and space of an important life-history trait. Estimating fitness-related traits within and among individuals in multiple populations through time using a trait-based approach is of fundamental interest in evolution and ecology, and our work can inform others wishing to partition out variation for similar questions. Size and growth rate are important components of life-history that need to be evaluated correctly and nuanced modelling may sometimes be necessary to understand their multifaceted effects on life-history, especially in heterogeneous environments. Excluding trait-based variation within species ignores the basic fact that particularly in fish, individuals feed at different trophic levels and are prey to different species throughout their ontogeny, related to their size (72). Interestingly, by integrating size-dependence into our models, we showed that there was more size-*independent* than size-dependent variation in growth. This suggests that variation in environmental conditions in space, time and space-time affects individuals across sizes similarly in this ecosystem.

## Data availability statement

Data and code are available on OSF: https://osf.io/nsuw3/?view_only=480c19c86a2a478da7bf8a6a2c8969b4.

## Acknowledgements

We would like to acknowledge the many field assistants and colleagues who helped collect the data throughout the years and the landowners allowing us to work on their land. We are especially grateful to Árni Einarsson and the Mývatn Research Station, Skúli Skúlason, Anett Relient, Mathias Lherbier and Kári H. Árnason. The long term monitoring of Arctic charr in lava caves is funded by the Icelandic Research Fund, RANNÍS (research grants 120227 and 162893). We would like to thank Ben Bolker and Timothée Bonnet for useful comments on our work. E.A.M. was supported by the Icelandic Research Fund, RANNÍS (162893) & a NERC research grant awarded to M.B.M. (NE/R011109/1). M.B.M. was supported by a University Research Fellowship from the Royal Society (London). C.A.L. & B.K.K. were supported by Hólar University, Iceland. All animals were captured, tagged and handled according to standard procedures validated by the Icelandic Food And Veterinary Authority through an animal care permit #1904736.

## Author Contributions and Statement on inclusion

C.A.L., B.K.K., K.R., M.M.F. & M.B.M. conceptualised the study; C.A.L., B.K.K., K.R., & E.A.M. collected the data; E.A.M. & M.B.M. analysed the data; E.A.M. wrote the original draft; All authors contributed critically to the manuscript and gave final approval for publication. Most authors have lived and worked in northern Iceland during their research careers, if not there now, and collected the data. Close collaboration within the country are maintained and work by other local researchers is cited.

