## Supplementary Materials for "Spatiotemporal variation in size-dependent growth rates in small isolated populations of Arctic charr (*Salvelinus alpinus*)"

### Supplementary Information for Spatiotemporal variation in growth rates in small isolated populations of Arctic charr (*Salvelinus alpinus*)

##### Model Fit

We generated data to test the fit of our model by randomly removing a single observation from ten randomly chosen individuals and re-running the model (example in Fig. 1). As with the main model four independent chains were run each time to check for model convergence. This was done 99 times giving us 990 individual values predicted by the model to compare to real data observations.

The vast majority of the deleted observations fell within the 95% CI of values predicted by the model ($\sim$ 93%; e.g., individual observations were within the grey bars in Fig. 1): 4.5% (45/990) were higher than the predicted value and 3.1% (31/990) were lower than the predicted value (95% CIs not overlapping the 1:1 line in Fig. 2). Furthermore, the observed and predicted estimates of growth based were almost identical (Figure 3). Although there could be a small visual trend in the residuals for a slight underestimation of the size of smaller fish (<50 mm; residuals were calculated as observed values - predicted values; Fig. 4a), the bulk of our data is from larger fish (95% range: 52-134 mm) and so this apparent trend is unlikely to have biased our results. Further, fish growth in general, and in these Arctic charr, is size-dependent so larger fish are growing less. Therefore, previous and subsequent sizes will also be more informative for the size of larger fish at each time point. Overall, the model is likely to struggle more with the smallest fish in this system due to less of information for these smaller sizes. This is supported by Pearson's correlation coefficient between the residuals and the predicted values ($r = -0.127$), which suggests that the model is generally closer to the observed value (the residual is smaller) as fish size increases. We can see in Fig. 4b that this is not biased in either a positive or negative direction, but is an increase in the accuracy of the predicted size.

The overall Pearson's correlation coefficient between the predicted and observed sizes was $r = 0.982$, suggesting a good fit, and the mean absolute residual was 2.85 mm (root mean squared error = 4.99 mm). There are several reasons why we do not expect there to be a "perfect" fit of our model to the data. Firstly, a perfectly fitted model for a system is often a sign of overfitting a model, and could particularly be the case in this Arctic charr system where we have multiple populations within one study. Secondly, the predicted values do not include all the practicalities of making real life observations, such as measurement error (although the model does include an estimate of this; see main manuscript methods equation 4). Finally, the predicted values are estimates of size on the mid-date of each sampling occasion, which contains two visits approximately two weeks apart. The observed data will be measured on earlier and/or later dates than this mid-date (taken into account in the main model; see main manuscript methods equation 4).

No patterns were seen in the residuals based on the temporal distance (in numbers of sampling occasions) to another observed size (Fig. 5). However, some individuals were only seen during one capture occasion, which could include multiple observations as each capture occasion contained two sampling periods (e.g., k=0 in Fig. 5a-d). In order to make sure this was not biasing our results we removed data from the dataset where there was observed data within the same sampling occasion as the predicted value (k=0) and re-ran our analyses. There was basically no change in Pearson's correlation coefficient between the predicted and observed sizes ($r = 0.990$), and the correlation between the residuals and the predicted values remained low ($r = 0.142$). Approximately $\sim$ 96% of the observations still fell within the 95% credible intervals (2.3% higher and 1.9% lower). Overall, these checks suggest that the data augmentation scheme is working well.

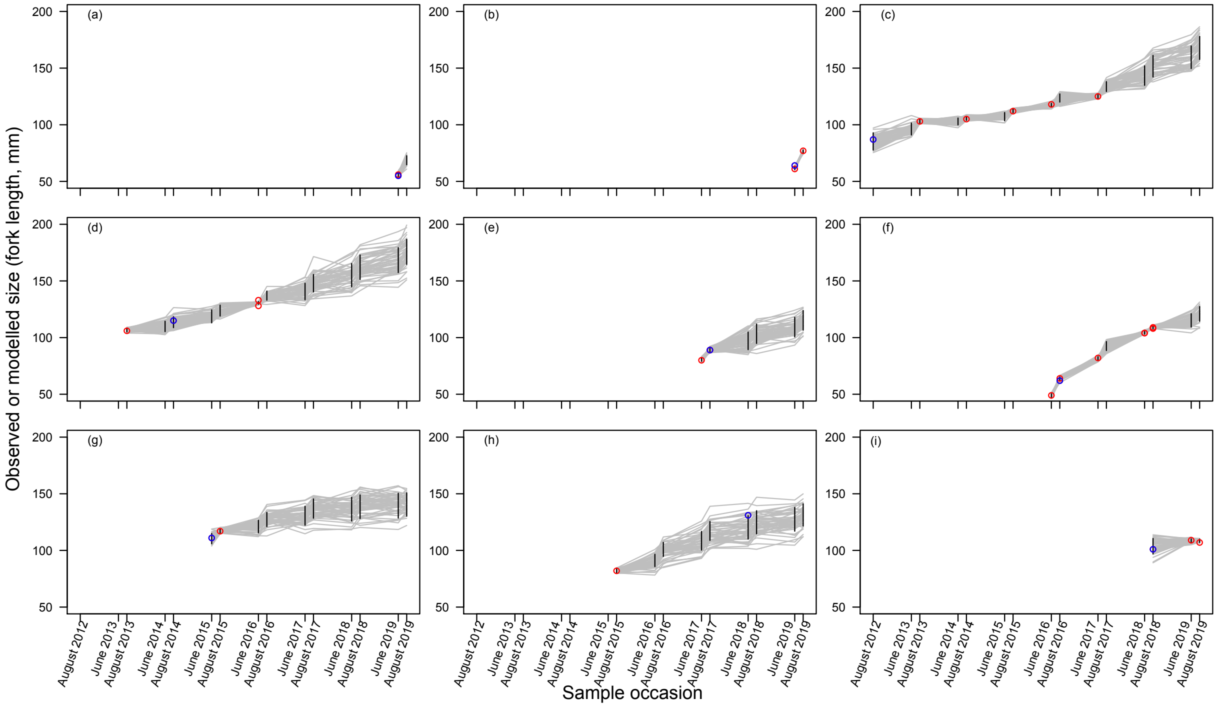

Figure 1: Examples of individual growth trajectories for a random selection of individuals with different observed capture histories. For each individual a single observation was randomly deleted. The red points are the observed data that is given to the model, and the blue points are the observations that were deleted and so not seen by the model. These individuals were traced in the model to obtain the modelled individual trajectories and the uncertainty around them. The different capture histories of the individuals shown are: (a), (b) and (c) individuals who were caught in the first ($k = 1$) and last capture occasions ($k = 15$); (d), (e) and (f) individuals that were caught twice with a large gap between (i.e. caught once and not seen for at least six $k$, then caught again); (g) and (h) individuals that were caught in the first capture occasion ($k = 1$) but never seen again; (i) an individual who was only caught twice in consecutive years. The red points are the observed sizes (fork length mm). The grey lines are modelled trajectories with the black vertical lines indicating the posterior distribution for size at each capture occasion where an individual was not observed.

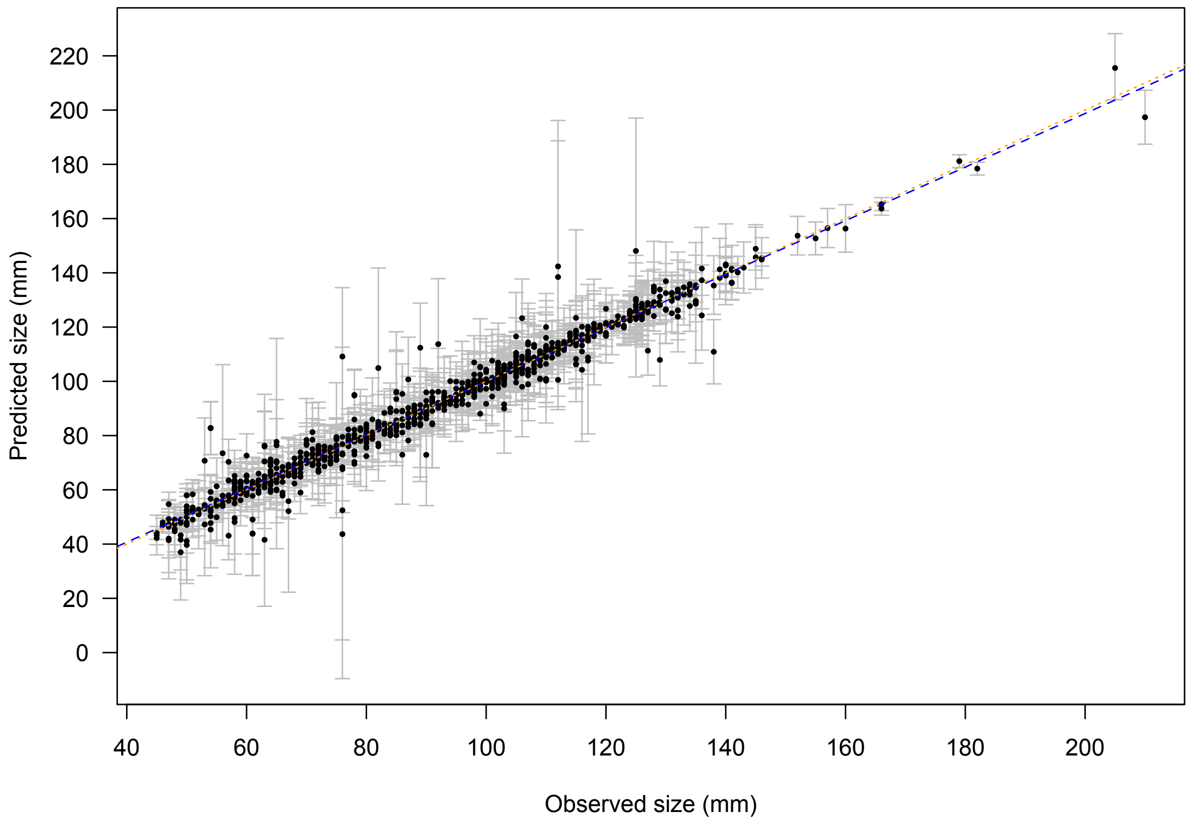

Figure 2: The model predicted sizes (mm) are shown on the y-axis and the real observed sizes (mm) that were deleted are on the x-axis. The blue dashed line is a simple linear regression between the two sets of values and the orange dashed line is the 1:1 line. These lines are almost identical. The 95% credible intervals are shown around the predicted values with grey bars.

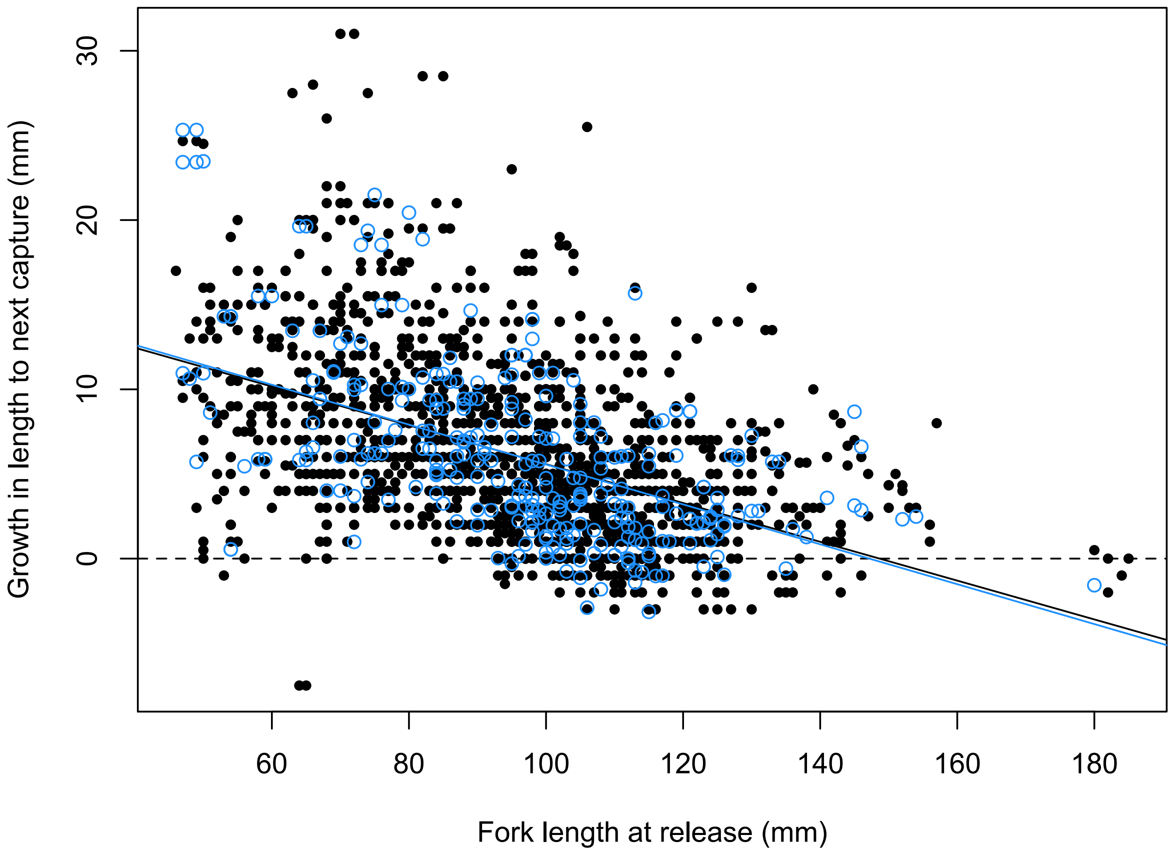

Figure 3: Fork length at release (mm) is shown on the x-axis and growth to the next capture occasion (mm) is shown on the y-axis. The observed (black points) and predicted (blue circles) growth is shown. The black and blue lines are simple linear regressions between the two sets of values, with black and blue representing observed and predicted growth, respectively. These lines are almost identical.

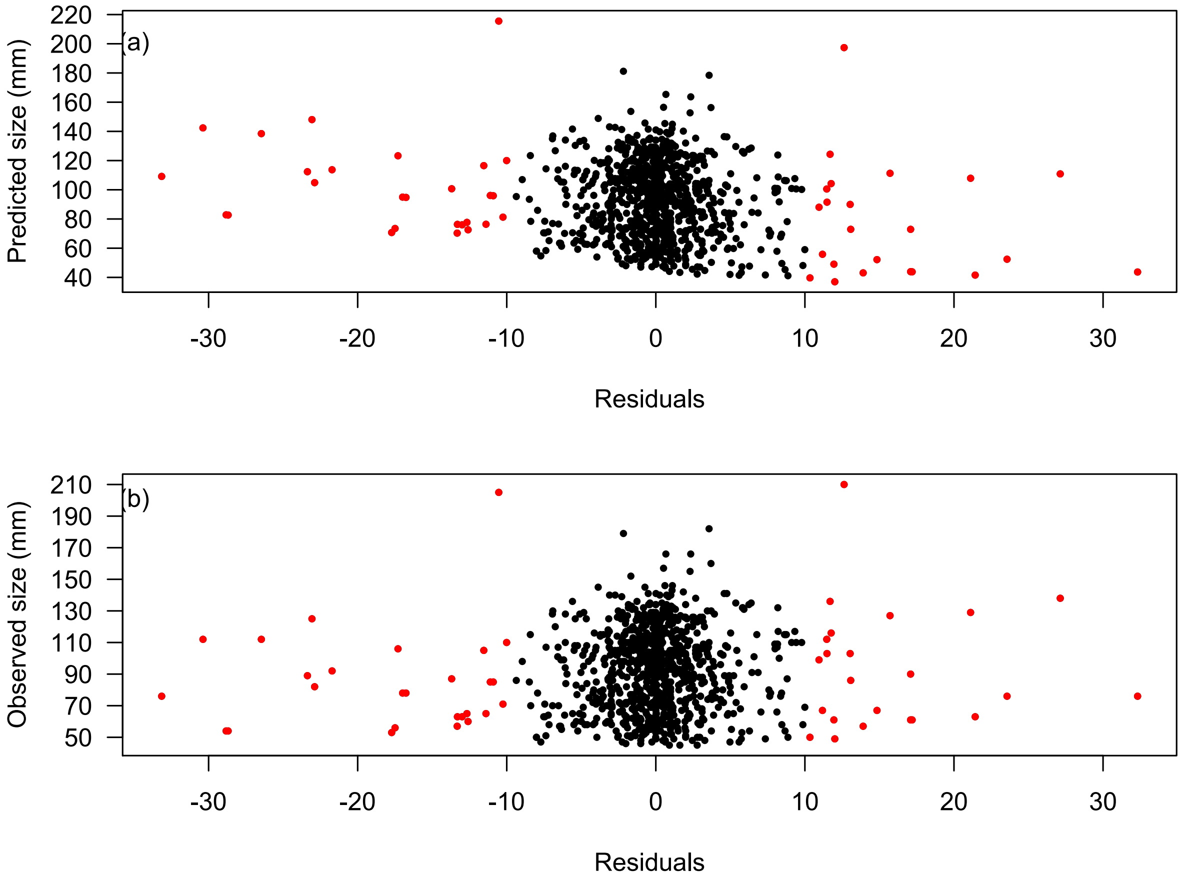

Figure 4: The residuals (observed - predicted) are shown on the x-axis. Figure (a) shows the residuals plotted against the values predicted by the model on the y-axis and figure (b) shows the residuals plotted against the observed data on the y-axis. Predictions that are greater than 10 mm from the observed value are highlighted in red.

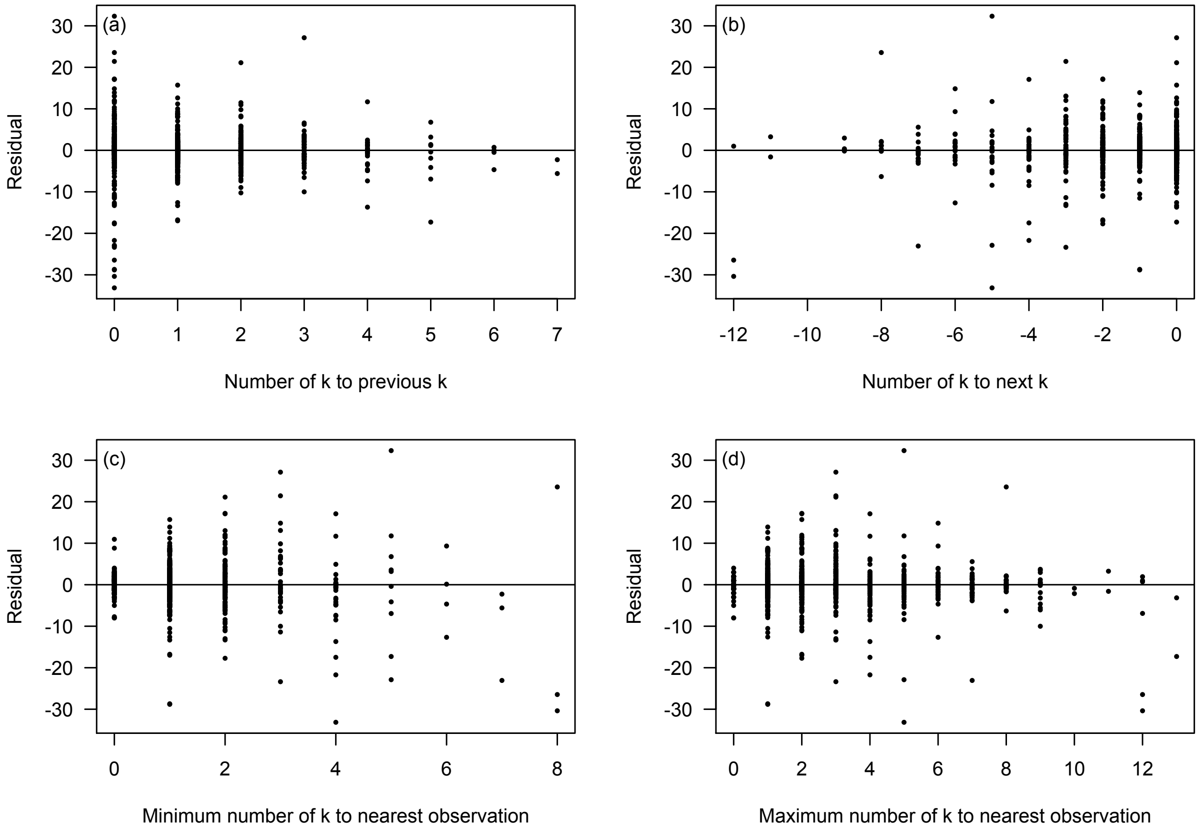

Figure 5: The residuals (observed - predicted) for size (mm; y-axes) are shown against: (a) the number of sampling occasions (k) between the observation and the previous observation; (b) the number of k between the observation and the next observation; (c) the minimum number of k between the deleted observed data and another observation; and (d) the maximum number of k between the deleted observed data and another observation. The horizontal line indicates a perfect predicted size relative to the observed size.

#### Additional figures

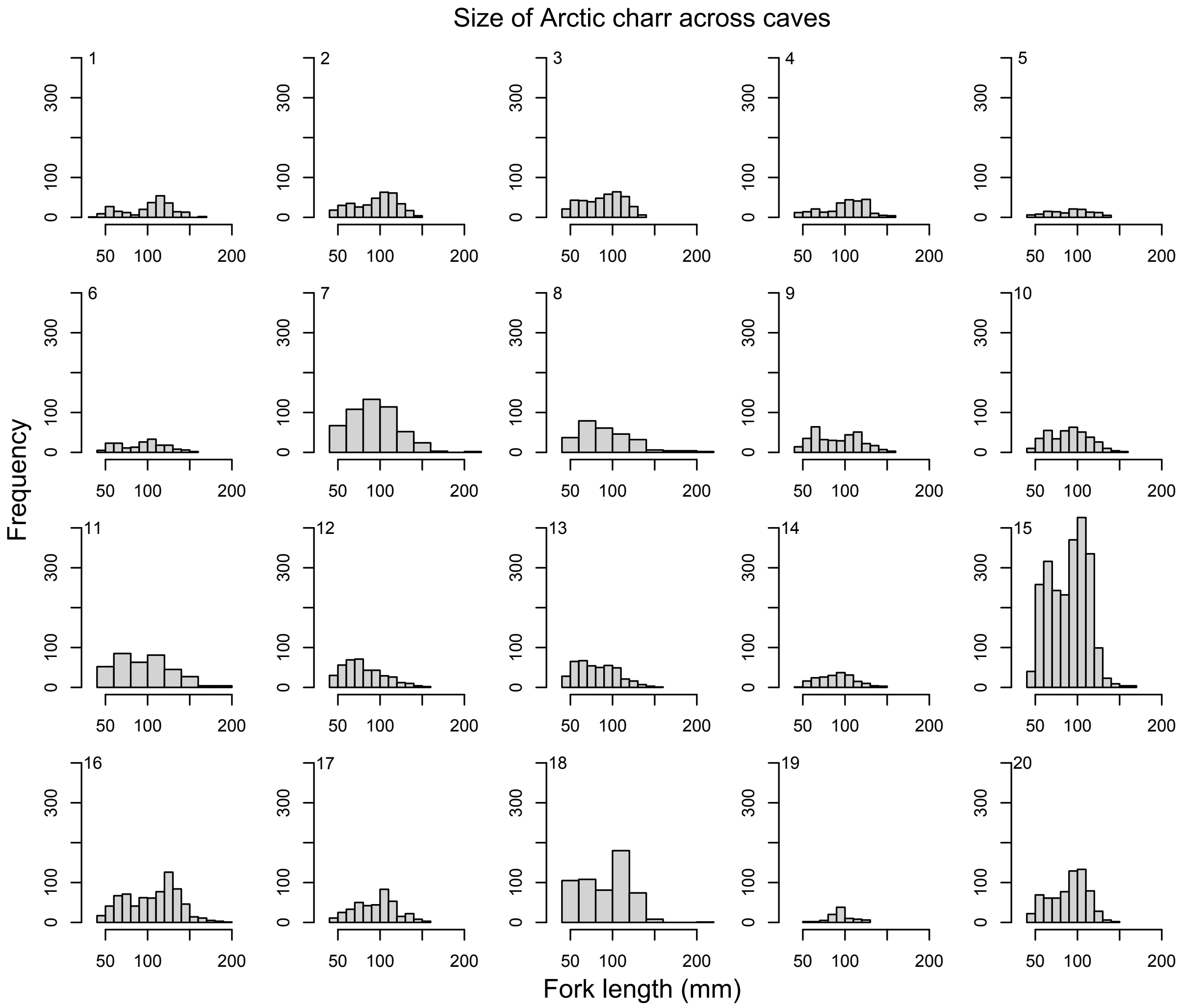

Figure 6: Raw size data across caves.

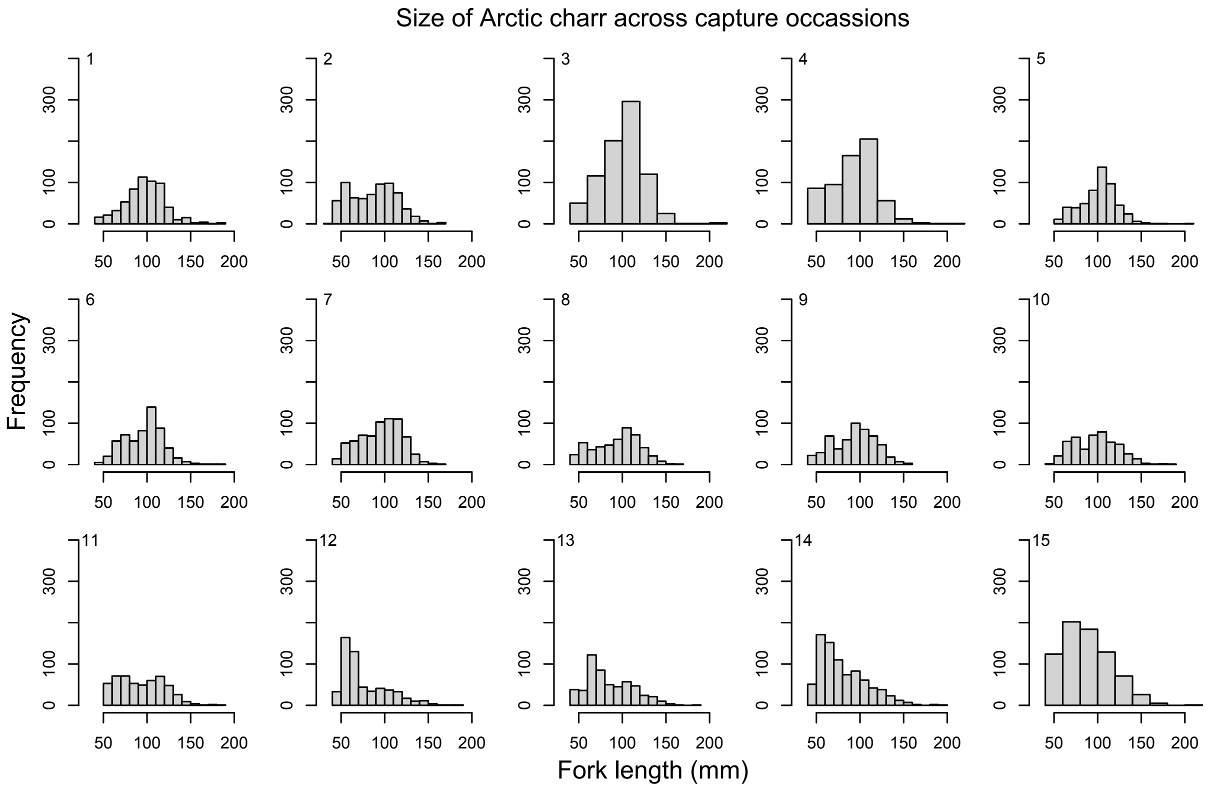

Figure 7: Raw size data across capture occasions.

#### Covariances between size-dependent and size-independent growth

Two of the covariances between the slopes (size-dependent growth) and intercepts (size-independent growth) of growth rate are negative and have credible intervals that do not overlap zero: the temporal and spatiotemporal estimates in winter (Table 1). Visualising this relationship the pattern appears to be stronger temporally (Figure 8). However, considering the magnitude of the estimates (perhaps clearer when looking at the point estimates in Figure 9 or thinking about the amount of variation present in the slopes), the changes in the slope fall within a very small range (-0.15 – 0.15 mmˆ2). A negative covariance would occur when the size-dependence (slope) decreases when there is an increase in the size-independent growth (intercepts), or in other words, when all sizes are experiencing higher growth rates there is less dependence of that growth on size.

Table 1: The covariance between the slopes (size-dependent growth) and intercepts (size-independent growth) in space, time and space-time in summer and winter from our model of growth rate in Arctic charr. The 95% credible intervals are shown in brackets.

| | Summer | Winter |
|---|---|---|
| Spatial | 0.070 [-0.545; 0.646] | 0.436 [-0.141; 0.472] |
| Temporal | 0.319 [-0.511; 0.920] | -0.682 [-0.978; -0.072] |
| Spatiotemporal | -0.088 [-0.537; 0.366] | -0.357 [-0.664; -0.020] |

Table 2: Overall estimates of coefficients from our model of monthly average growth rates in Arctic charr. The 95% credible intervals are shown in brackets.

| | Summer | Winter |
|---|---|---|
| Size-independent estimates | 3.33 [2.38; 4.26] | 0.81 [0.51; 1.12] |
| Size-dependent estimates | -0.04 [-0.06; -0.03] | -0.02 [-0.02; -0.01] |
| Size-independent estimates associate with temperature | 0.405 [0.10; 0.705] | -0.014 [-0.096; 0.07] |
| Size-dependent estimates associated with temperature | 0 [0; 0.01] | 0 [0; 0] |

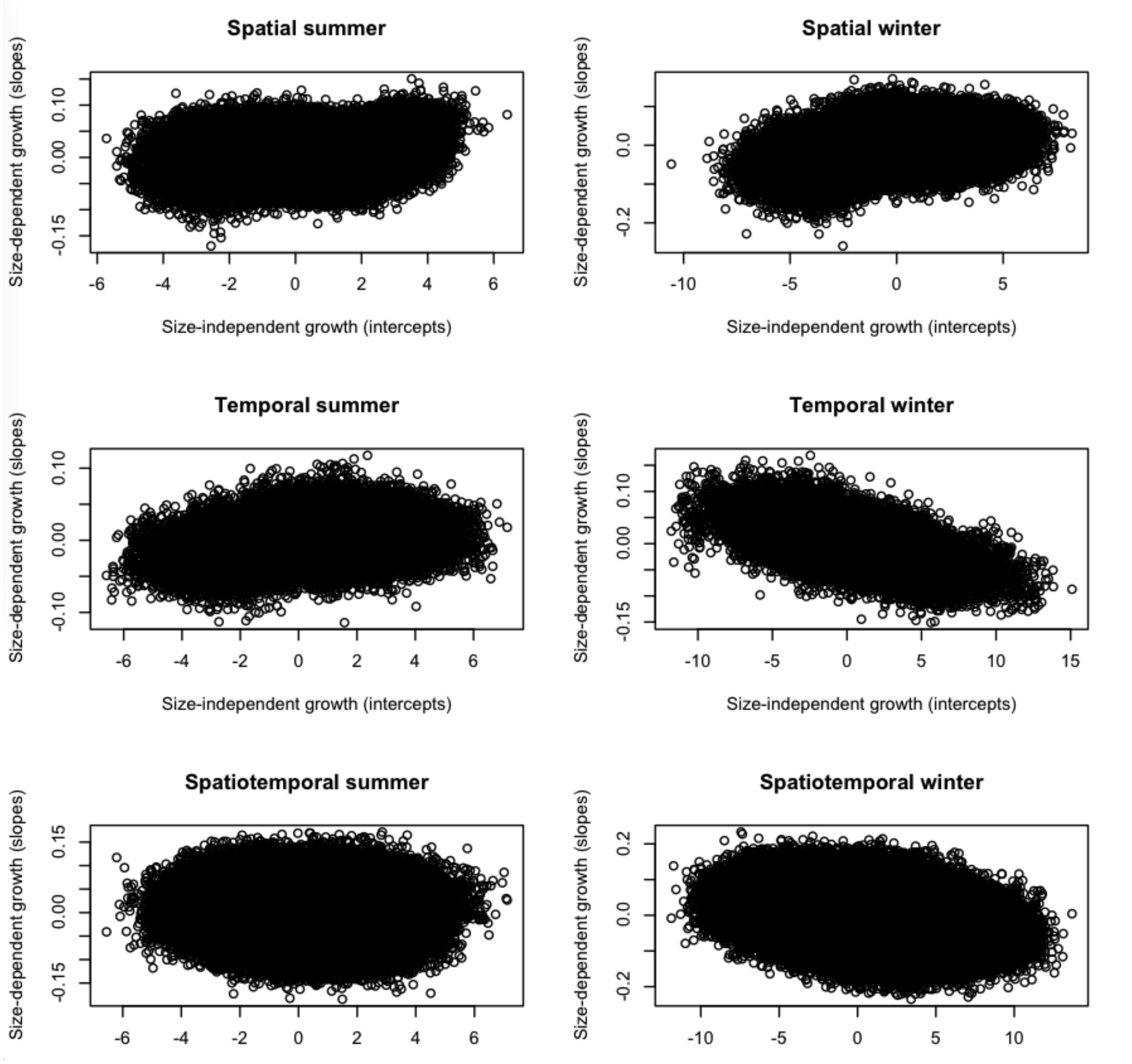

Figure 8: All estimates of slopes (size-dependent growth) and intercepts (size-independent growth) in space, time and space-time for summer and winter intervals of growth.

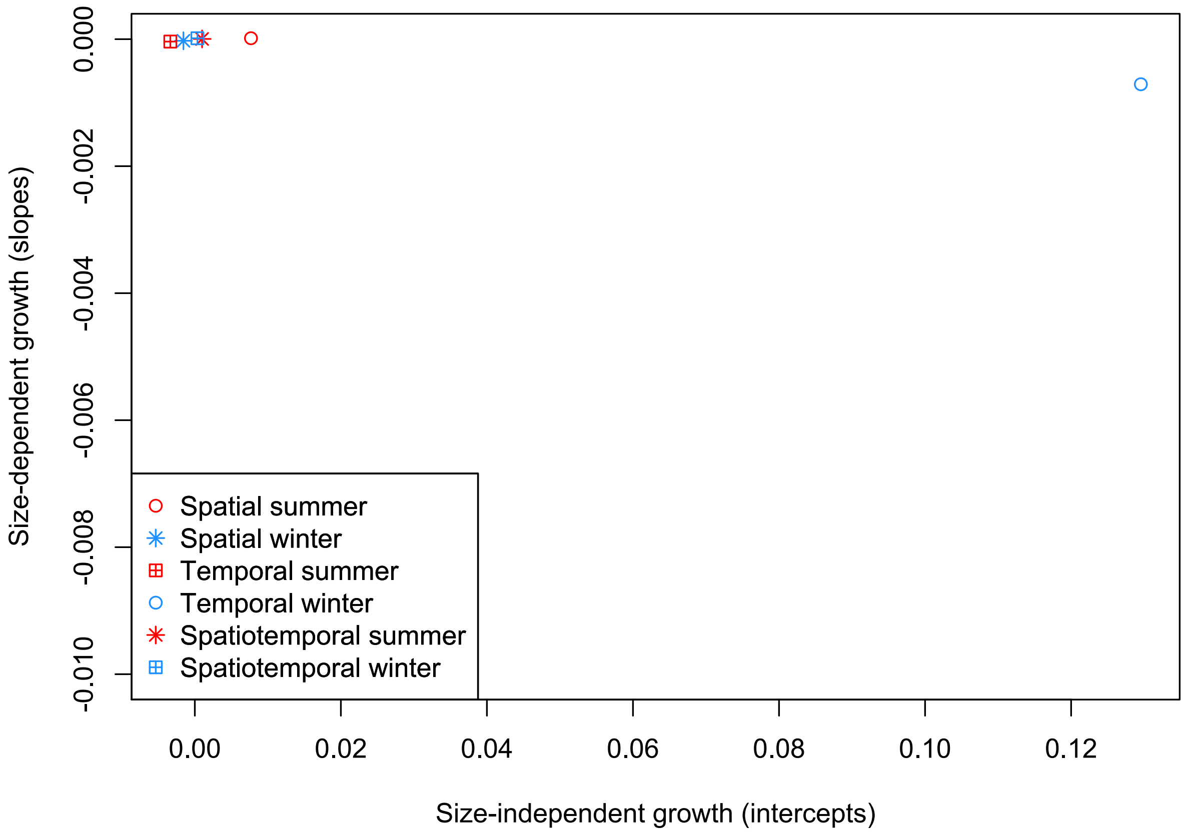

Figure 9: Average slopes (size-dependent growth) and intercepts (size-independent growth) in space, time and space-time for summer and winter intervals of growth.

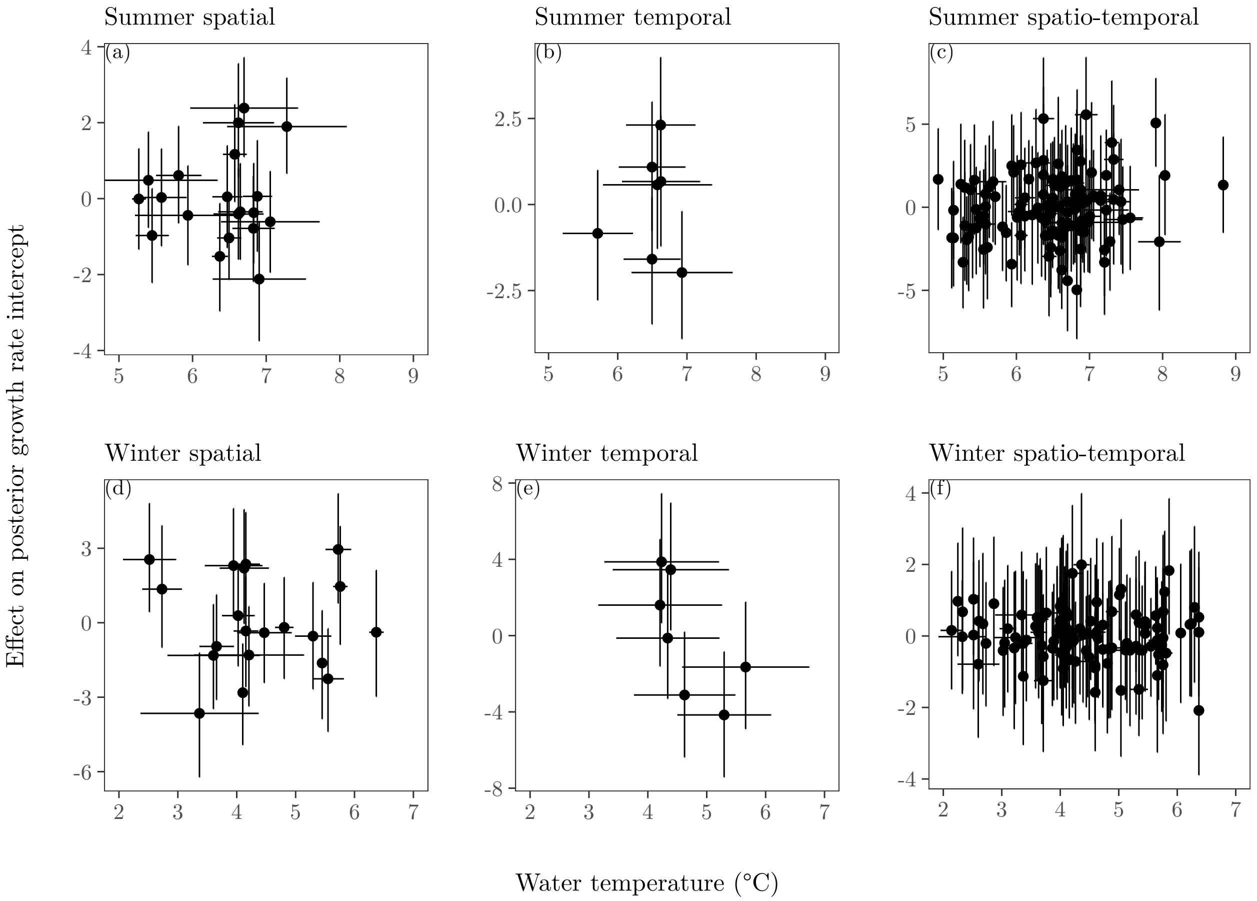

Figure 10: Spatial, temporal and spatial-temporal variation in water temperature (°C; x-axis) and the spatial, temporal, and spatial-temporal effects on the posterior distribution of the growth intercept (y-axis) for summer (top row; a-c) and winter (bottom row; d-f). The error bars represent the standard deviation for water temperature.

Table 3: Variation in mean estimates of monthly growth rate through time, space and space-time shown for summer and winter seperately in Arctic charr. The 95% credible intervals are shown in brackets.

| | | Summer | Winter |
|---|---|---|---|
| Size-independent estimates | Space | 0.004 [-1.269;1.422] | 0.000 [-0.405;0.392] |
| | Time | -0.002 [-1.586;1.635] | 0.013 [-0.556;0.597] |
| | Space-time | 0.000[-2.368;2.629] | 0.000 [-0.223;0.228] |
| Size-dependent estimates | Space | 0.000 [-0.005;0.005] | 0.000 [-0.060;0.063] |
| | Time | 0.000 [-0.004;0.004] | -0.001 [-0.061;0.062] |
| | Space-time | 0.000 [-0.008;0.008] | 0.000 [-0.057;0.057] |

#### Asymptotic size thought experiment

An old fish will experience not a single growth function, but a number of growth functions, due to temporal variation. Variation in $L_\infty$ does not correspond to any variation that can be observed among the sizes of old fish in reality, unless reality happens to be that there is very little temporal, or time/space, heterogeneity in growth. What can be, and is, calculated here is a slightly tricky thought experiment where snapshots of growth functions are imposed on a long-lived organism. The expectation of asymptotic size in space, time and space-time for the two seasons was estimated using

$$\mathbb{E}[L_{\infty(\alpha,\beta)}] = L_{ref} - \frac{\alpha}{\beta} - \frac{\alpha\sigma_\beta^2}{\beta^3} + \frac{\sigma_{(\alpha,\beta)}}{\beta^2} \tag{1}$$

where $\alpha$ is the intercept (size-independent growth), $\beta$ is the slope (size-dependent growth), $L_{ref}$ is 92 mm (the reference size used in the model), $\sigma_\beta^2$ is the total variation in slopes across space, time and space-time in summer and winter, and $\sigma_{(\alpha,\beta)}$ is the covariance between the intercept and slope for each. This is an approximation based on the von Bertalanffy growth function (Von Bertalanffy 1957) following Appendix 1 in Lynch and Walsh (1998). Similarly, the variation in asymptotic size is approximated using

$$\sigma^2[L_{\infty(\alpha,\beta)}] = \frac{\sigma_\beta^2\alpha^2}{\beta^4} + \frac{\sigma_\alpha^2}{\beta^2} - \frac{2\alpha\sigma_{(\alpha,\beta)}}{\beta^3}. \tag{2}$$

As mentioned above, none of the fish are experiencing the same conditions across space, time and space-time in summer and winter, and individuals will likely grow when conditions are good but not shrink in poor conditions. Therefore, although these approximations are the one way to assess asymptotic size with these data, there are two reasons why we do not expect these approximations to be accurate. Firstly, we are extrapolating out of the growth model to sizes of fish that we do not have data for (and may not exist). Secondly, this is a linear approximation for a function that is likely to be non-linear. To go even further, we used Monte Carlo sampling to estimate of the expectations as a comparison. In this sampling case we are still predicting out of the size range for a linear function. Overall, the first order approximation looks like it could be an underestimation of the asymptotic size (Supplementary Table 4).

Table 4: Analytical and Monte Carlo approximations of the expectations of asymptotic size in these Arctic charr, in summer and winter, are shown in the top half of the table. The analytical approximations of the variances in the asymptotic size in these Arctic charr, in summer and winter, and through space, time and space-time, are shown in the bottom half of the table. 95% confidence intervals are shown in brackets.

| Expectations | Summer | Winter |
|---|---|---|
| Analytic approximation | 139.784 [125.119; 238.401] | 99.558 [112.343; 136.853] |
| Monte Carlo approximation | 265.547 [-300.37; 776.113] | 136.277 [68.341; 234.586] |
| Variances | Summer | Winter |
| Space | 867.054 [382.553; 8191.232] | 476.117 [-5.237; 527.985] |
| Time | 1895.166 [32.704; $1.2038701 \times 10^4$] | 380.425 [-404.674; 662.047] |
| Space-time | 699.140 [668.968; 7431.24] | 476.117 [-52.462; 418.608] |

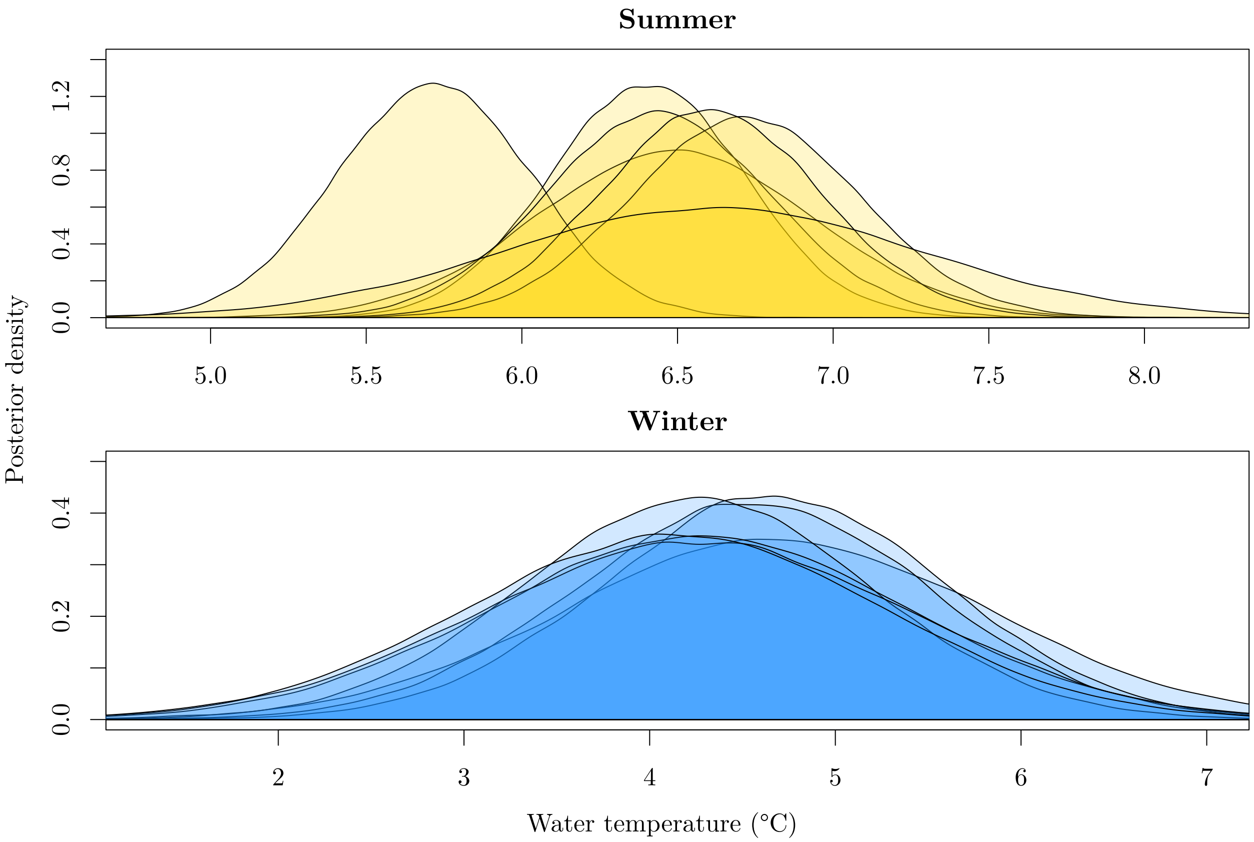

Figure 11: Posterior distributions of water temperature (°C) across years for summer (top row) and winter (bottom row) as estimated by the model $T_{j,t}$ in Equation 1. Note that both temperatures and range of temperatures on the x-axes differ for the two seasons as water temperature is lower during the winter months.

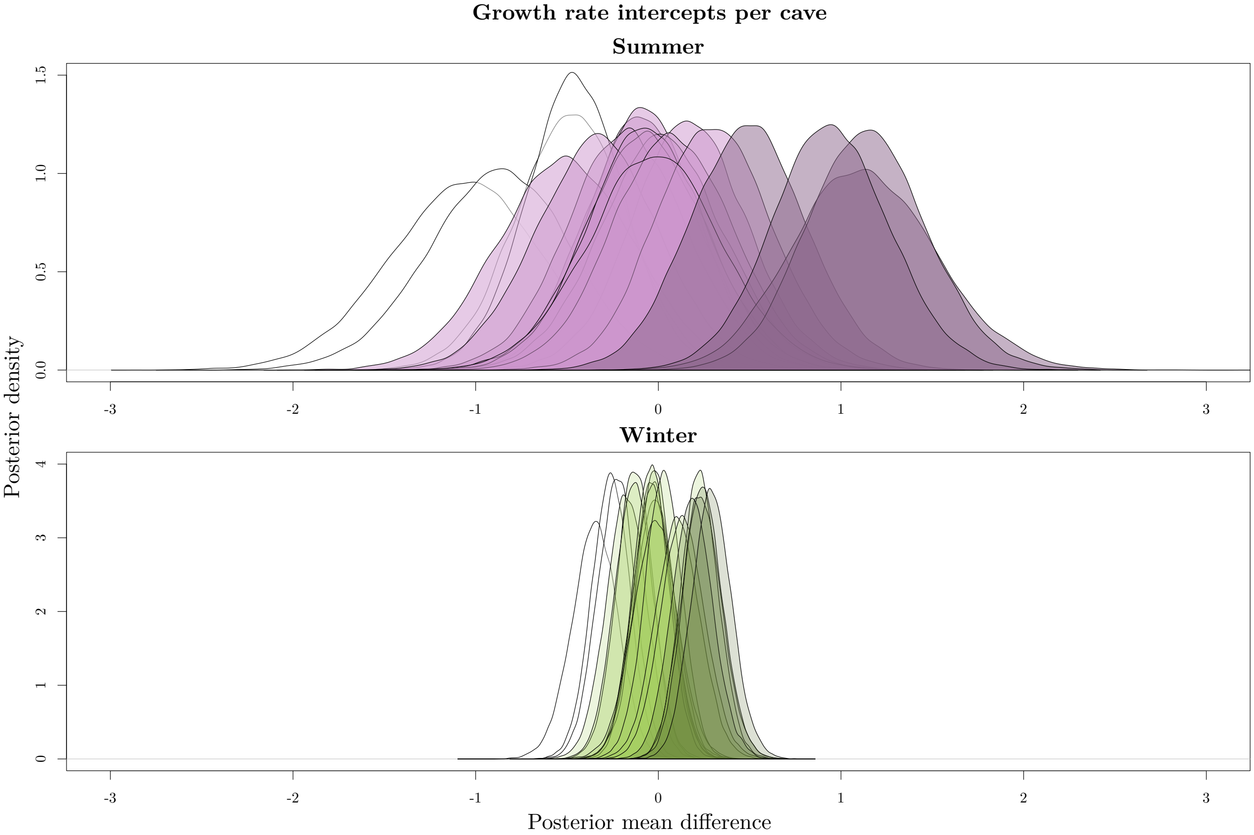

Figure 12: Posterior distributions of the cave effect on growth intercepts in the summer and winter using all capture occasions. The growth rate is standardised to mm/month. Caves with lower than average growth are indicated by white and those with higher than average growth are indicated with a darker shade. In both seasons caves 6, 7 and 25 had lower than average growth and caves 26, 21 and 22 had higher than average growth. In summer cave 17 also had lower than average growth and in winter cave 19 had higher than average growth.

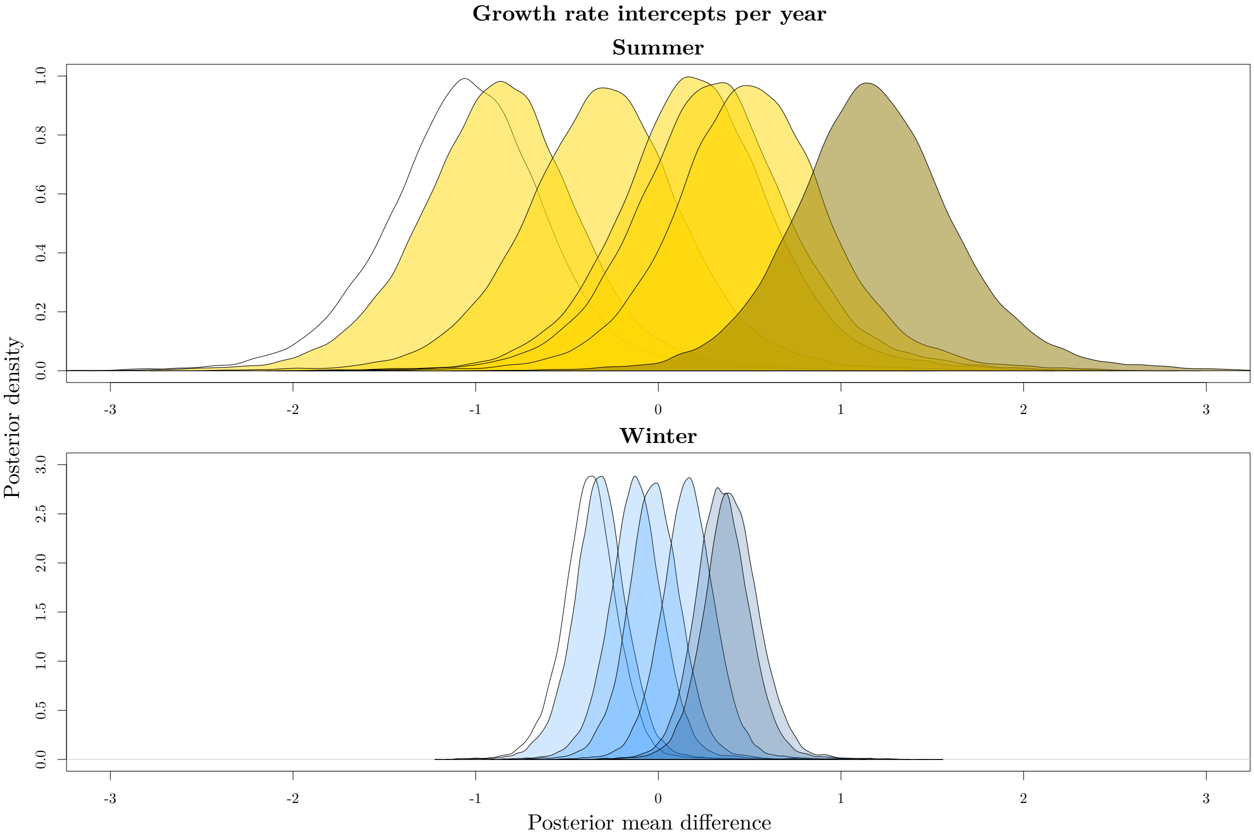

Figure 13: Posterior distributions of the year effect on growth intercepts in the summer and winter using all caves. The growth rate is standardised to mm/month. Years with lower than average growth are indicated by white and those with higher than average growth are indicated with a darker shade. In both seasons 2014 had lower than average growth and 2017 had higher than average growth. In addition, in winter 2018 had higher than average growth. Growth was almost lower than average in both seasons for 2013 and 2015.

#### Variation in size models

To estimate the variation in fork lengths (size) mixed-effects models were run in `R` package `MCMCglmm` (Hadfield 2010). These models were run seperately for summer and winter and included random effects of cave and year. The code is shown below.

```
library(MCMCglmm)
# Get data
data <- read.csv("model_data.csv")
data <- data[,-1]
data$fl <- as.numeric(data$fl)

# Summer and winter separately
# Summer is the odd k's
data_sum <- subset(data[which(data$k %in% c(3,5,7,9,11,13,15)),])
# 4399 observations
prior_sum <- list(R=list(V=1,nu=0.002),G=list(G1=list(V=1,nu=2),G2=list(V=1,nu=2)))
model_sum <- MCMCglmm(fl ~ 1, random = ~cave_ori + year_ori, family = "gaussian",
                      nitt=110000, burnin=10000, thin=10, verbose=FALSE,
                      prior=prior_sum, data=data_sum)
# Check convergence
plot(model_sum)
# Autocorrelation
autocorr.diag(model_sum$VCV)
# Effective sample sizes
summary(model_sum)

# Winter is the even k's
data_win <- subset(data[which(data$k %in% c(2,4,6,8,10,12,14)),])
# 4258 observations
prior_win <- list(R=list(V=1,nu=0.002),G=list(G1=list(V=1,nu=2),G2=list(V=1,nu=2)))
model_win <- MCMCglmm(fl ~ 1, random = ~cave_ori + year_ori, family = "gaussian",
                      nitt=110000, burnin=10000, thin=10, verbose=FALSE,
```

```r
                        prior=prior_win, data=data_win)
# Check convergence
plot(model_win)
# Autocorrelation
autocorr.diag(model_win$VCV)
# Effective sample sizes
summary(model_win)
```
